# Genome editing of the *vermilion* locus generates a visible eye color marker for *Oncopeltus fasciatus*

**DOI:** 10.1101/2022.12.20.521233

**Authors:** Katie Reding, Minh Lê, Leslie Pick

## Abstract

Insects display a vast array of eye and body colors. Genes encoding products involved in biosynthesis and deposition of pigments are ideal genetic markers, contributing, for example, to the power of *Drosophila* genetics. *Oncopeltus fasciatus* is an emerging model for hemimetabolous insects, a member of the piercing-sucking feeding order Hemiptera, that includes pests and disease vectors. To identify candidate visible markers for *O. fasciatus*, we used parental and nymphal RNAi to identify genes that altered eye or body color while having no deleterious effects on viability. We selected *Of-vermilion* for CRISPR/Cas9 genome editing, generating three independent loss-of-function mutant lines. These studies mapped *Of-vermilion* to the X-chromosome, the first assignment of a gene to a chromosome in this species. *Of-vermilion* homozygotes have bright red, rather than black, eyes and are fully viable and fertile. We used these mutants to verify a role for *Of-xdh1*, ortholog of *Drosophila rosy*, in contributing to red pigmentation after RNAi. Rather than wild-type-like red bodies, bugs lacking both *vermilion* and *xdh1* have bright yellow bodies, suggesting that ommochromes and pteridines contribute to *O. fasciatus* body color. Our studies generated the first gene-based visible marker for *O. fasciatus* and expanded the genetic toolkit for this model system.

## Introduction

Recent years have witnessed an explosion of genome sequencing and gene annotation for representative species across the tree of life. Understanding the roles these genes play in generating, maintaining, or even limiting biodiversity will require approaches to assess gene function at key branches of the tree. Insects are excellent subjects of investigation because of the large number of species, their diverse habitats, physiology, behaviors and morphologies. Additionally, many can be reared in lab settings to allow for molecular genetic analyses. Most studies of insects to date have focused on the holometabolous insects, including beetles, butterflies and flies, with *Drosophila melanogaster* among the most powerful genetic model systems. As a close outgroup to holometabolous insects, the order Hemiptera comprises >80,000 described species^1^ that share a piercing-sucking mode of feeding but which are otherwise enormously diverse (reviewed in^2,3^). This order includes agricultural pests such as aphids and harlequin bugs, nuisance pests of humans such as bed bugs and stink bugs, and vectors of serious human disease, such as kissing bugs, which transmit Chagas disease^4,5^. The milkweed bug *Oncopeltus fasciatus (Of)* is a promising model system for Hemiptera: a genome sequence is available and multiple tools for molecular genetic analyses have been developed, including RNA interference (RNAi) and CRISPR/Cas9 mutagenesis^6–9^. However, lacking for this species and Hemiptera more broadly are visible markers that are needed to pursue more rigorous genetic studies.

Pigmentation has long fascinated researchers and has been the target of genetic approaches because it varies extensively, even within species, and is simple and direct to assay, being readily visible externally. Mutations in genes that cause variation in body or eye color often do not impact organismal viability, allowing for isolation and maintenance of mutant lines. This was exploited in the early 1900s to launch the field of *Drosophila* genetics^10,11^. At that time, studies of eye color mutants from several different insect species contributed to the “one gene-one enzyme hypothesis” and established an ordered biochemical pathway involved in synthesis of insect pigments^12–17^. Body color in some insects is based on melanin (reviewed in^18^). However, the brightly colored eye and body colors of many insects are comprised of two other pigment types: ommochromes (named from the Greek *omma* - eye, and *khroma* - color) which derive from tryptophan; and pteridines (named for the Greek *pteron* – wing, based on their first identification from the orange, yellow and white colors in the wings of Pieridae butterflies (see^19^), which are derived from GTP. Overall, ommochrome pigments are in the brown range, with mutations generating red- or white-eyed insects, while pteridines range from bright red to yellow, with mutations often leading to loss of body pigmentation and brown eyes (reviewed in^20,21^). Biosynthetic pathways leading to ommochrome pigments include products of the *karmoisin (kar)*, *vermilion* (*v*), *cinnabar* (*cn*) and *cardinal* (*cd*) genes^22–24^ (recently reviewed^25^), while biosynthesis of pteridine pigments requires gene products of *punch (pu)* and *purple (pr)*, and includes additional enzymatic steps, such as the product of *rosy (ry)* which encodes xanthine dehydrogenase, to generate a variety of colors^19,26^ (recent review^27^). For each pathway, specific transporters are required for pigment deposition^28^. The genes *white (w), scarlet (st),* and *brown (bw)* are closely related and comprise a subfamily of the ABC-transporter-encoding genes, with White forming a heterodimer with Brown for pteridine precursor transport, and with Scarlet for ommochrome precursor transport^29–34^. In *Bombyx mori,* another member of this subfamily, *ok,* was identified and found to be most closely related to *Bm-brown; Bm-ok* mutants display defects in uric acid deposition in the larval epidermis resulting in a translucent skin^35^. Although no *ok* ortholog was found in the *Drosophila* genome, orthologs have been identified in many other insect genomes.

Hemiptera display diverse colors and patterns that include contributions from melanin (reviewed in^18,36^) in addition to ommochromes and pteridines. Roles for these latter pigment types in color variation have been investigated recently for a number of species within this order. Brent & Hull^37^ combined gene expression analysis and RNAi to document the roles of five ommochrome pathway genes in eye color development in *Lygus hesperus (Lh)*. Knockdown of *Lh*-*bw* suggested that the pteridine pathway contributes to eye color while *w* knock down impacted both eyes and body, with phenotypes including smaller and paler bodies and soft cuticles. Subsequent CRISPR/Cas9 mutagenesis in this species showed that the ommochrome pathway components *Lh*-*cn* and *Lh*-*cd* are both involved in eye pigmentation, with *Lh*-*cn* mutants displaying sex-specific phenotypic differences in adults^38^. In the brown planthopper, *Nilaparvata lugens*, CRISPR/Cas9 genome editing suggested that *Nl*-*w* and ommochrome pathway member *Nl*-*cn* are necessary for wild type eye color^39^. Orthologs of ABC transporter genes *w*, *st* and *ok* as well as ommochrome pathway member *cn* were identified in the kissing bug *Rhodnius prolixus (Rpro)*, and their roles in pigmentation tested with RNAi^40^. Interestingly, nymphal RNAi knockdown of *Rpro-cn* resulted in animals with reddish, rather than black, ommatidia, but parental RNAi caused lethality post-blood meal in first instars. This suggests the *Rpro-cn* gene is required for functions other than eye color, making it a poor candidate for a visible marker gene. Transporter knockdown revealed a role for *Rpro-st* in ommochrome deposition in eyes, with newly added ommatidia displaying bright red color similar to that seen after *Rpro-cn* nymphal RNAi, and for *Rpro-w*, which also impacted viability, but no change in eye color was observed after knockdown of either *ok* ortholog. However, subtle changes in body pigmentation were noted after knockdown of *Rpro-st* or either of the two *Rpro-ok* paralogs, suggesting roles for both types of pigment in body color. Results from an extensive study of pigmentation-related genes in several water strider species^41^ suggested that ommochromes do not contribute to body color but are utilized for eye color in these species, with parental RNAi targeting either *cn* or *st* in *Limnogonus franciscanus* producing embryos with bright red eyes. Pteridines, on the other hand, appear to contribute to both eye and body pigmentation, with pRNAi of several different genes in that pathway producing loss of body coloration and brown eyes. Orthologs of *ry* were examined in several species where it is expressed and functions in both body and eye pigmentation.

*O. fasciatus* was one of the first hemipterans for which genetic approaches were taken to study pigmentation^42–44^.. Through an EMS mutagenesis screen, Lawrence and coworkers identified genes impacting pteridine and ommochrome pathways: *white body* (*wb*) resulted from a loss of specific pteridines such as erythropterin, the levels of which were found to coincide with the reddening of the body during embryogenesis^45^. Conversely, *red eye* (*re*) mutants had bright red eyes and wild type pteridines, and thus resulted from mutation in ommochrome synthesis. With both pteridine and ommochrome pathways disrupted, *wb/wb; re/re* mutants displayed loss of both red and brown pigments in the eye, resulting in white eyes and white bodies. In one study, Shelton and Lawrence (1974) made use of these mutants to show that ommatidia added as the compound eye grows through successive molts are recruited from cells of nearby epidermis^46^. After transplanting eye-proximal epidermal tissue from wild type nymphs into the corresponding region of *wb/wb; re/re* mutants, they observed mosaic eyes in the adult, mostly white eyes with clusters of wild type ommatidia at the eye margin closest to the graft, showing that these newer ommatidia had formed from transplanted epidermal tissue. The first study of these pathways using RNAi to target candidate genes in *O. fasciatus* was carried out by Liu^47^. Separate analyses of ommochrome pathway genes *v*, *cn* and *st*, and pteridine pathway genes *bw*, *pu*, *pr*, as well as *w* suggested roles for both pigment types in *O. fasciatus* coloration. Both parental and nymphal RNAi implicated pteridines in eye and body coloration, while nymphal RNAi knockdowns of ommochrome pathway genes generated animals with bright red eye margins but wild type body coloration, similar to the *re* mutant of Lawrence. Recently, RNAi knockdown demonstrated a role for *Of*-*red Malpighian Tubules* (*red*) in eye color, while also contributing somewhat to body pigmentation. This gene likely encodes a protein involved in intracellular pigment trafficking and, accordingly, knockdown impacted both ommochrome and pteridine pathways^48^.

In our first attempt to generate a visible eye color marker for *O. fasciatus*, we chose to target *Of*-*w*, given its historic use as a visible marker^49^. While that approach established CRISPR/Cas9 mutagenesis for *O. fasciatus*, we were surprised to find lethality in animals homozygous for *white* null alleles^9^, although *w* had been shown to be required for viability in the moth *Helicoverpa armigera*^50^ and more recently in *L. hesperus*^38^. Here, we used RNAi to assess roles of selected genes encoding candidate visible markers, testing for detrimental consequences of gene knockdown before proceeding to genome editing. Our experiments verified the role of *Of-v* in eye color^47,48^ and identified a novel role of the *Of*-*xdh*/*ry* gene in body color. We generated three independent *Of-v* null mutations, each of which has been maintained in lab colony for multiple generations without discernable impact on development. Thus, *Of-v* is an ideal visible marker for transgenesis, co-CRISPR, and other genetic tools, adding to the range of genetic studies feasible for this insect.

## Results

### *RNAi targeting* ok *paralogs fails to reveal clear role in pigmentation*

We first investigated two components of the pteridine pigment pathway: two previously identified *ok* orthologs in the *O. fasciatus* genome^9^. As ABC transporter-encoding genes closely related to *brown*, which is required for pteridine pigment transport in many insects (Introduction), the two *O. fasciatus ok* paralogs are good candidates for pteridine transport in this species. To evaluate the roles of these paralogs in *O. fasciatus* pigmentation, we designed two non-overlapping dsRNAs per gene. Following *Of-ok1* pRNAi, no difference in eye or body color in 1^st^ or 2^nd^ instar nymphs was observed compared to *gfp* controls (Fig. 1 A-B). Following *Of-ok2* pRNAi, very few embryos hatched despite appearing fully developed. Only 0.87% of embryos hatched following pRNAi with dsRNA-A (n=344), and 4.96% hatched after pRNAi using dsRNA-B (n=262), compared with an average hatch rate of 91.52% (n=544 across four replicates) for *gfp* controls (Table 1). Co-injection of both *Of-ok1* and *Of-ok2* dsRNA produced intermediate hatch rates (Table 1), but no change in coloration of hatched nymphs was observed. Furthermore, nRNAi knockdown of *Of-ok1* on 4^th^ instar nymph yielded no discernible change in eye or body coloration in either 5^th^ instar nymphs or adults (Fig. 1G) compared to *gfp* dsRNA-injected controls (Fig. 1F). In sum, despite being strong candidates for involvement in pteridine pigment transport in *O. fasciatus*, we found no evidence that either would serve as a useful visible marker gene for this species.

**Table 1.** Summary of hatch rates observed in pRNAi experiments.

| Gene(s) | dsRNA | dsRNA concentration (uM) | Analyzed (n) | # Hatch | Hatch rate (%) |
| --- | --- | --- | --- | --- | --- |
| *gfp* | A | 5 | 124 | 111 | 89.52 |
| *gfp* | A | 5 | 156 | 139 | 89.10 |
| *gfp* | A | 5 | 142 | 137 | 96.48 |
| *gfp* | A | 5 | 122 | 111 | 90.98 |
| *Of-ok1* | A | 5 | 262 | 240 | 91.60 |
| *Of-ok1* | B | 5 | 236 | 236 | 100.00 |
| *Of-ok2* | A | 5 | 344 | 3 | 0.87 |
| *Of-ok2* | B | 5 | 262 | 13 | 4.96 |
| *Of-ok1* + *Of-ok2* | A | 2.5 each | 327 | 155 | 47.40 |
| *Of-ok1* + *Of-ok2* | B | 5 each | 196 | 114 | 58.16 |
| *Of-st* | A | 5 | 188 | 179 | 95.21 |
| *Of-st* | B | 5 | 100 | 90 | 90.00 |
| *Of-v* | A | 5 | 257 | 246 | 95.72 |
| *Of-xdh1* | A | 5 | 190 | 184 | 96.84 |
| *Of-xdh1* | B | 5 | 272 | 199 | 73.16 |
| *Of-xdh2* | A | 5 | 208 | 182 | 87.50 |
| *Of-xdh2* | B | 5 | 189 | 188 | 99.47 |
| *Of-xdh1* + *Of-xdh2* | A | 2.5 each | 127 | 113 | 88.98 |
| *Of-xdh1* + *Of-xdh2* | B | 5 each | 87 | 76 | 87.36 |
| *gfp* ($v^3$ mut) | A | 5 | 255 | 255 | 100.00 |
| *Of-xdh1* ($v^3$ mut) | B | 5 | 212 | 204 | 96.23 |

**Figure 1.**
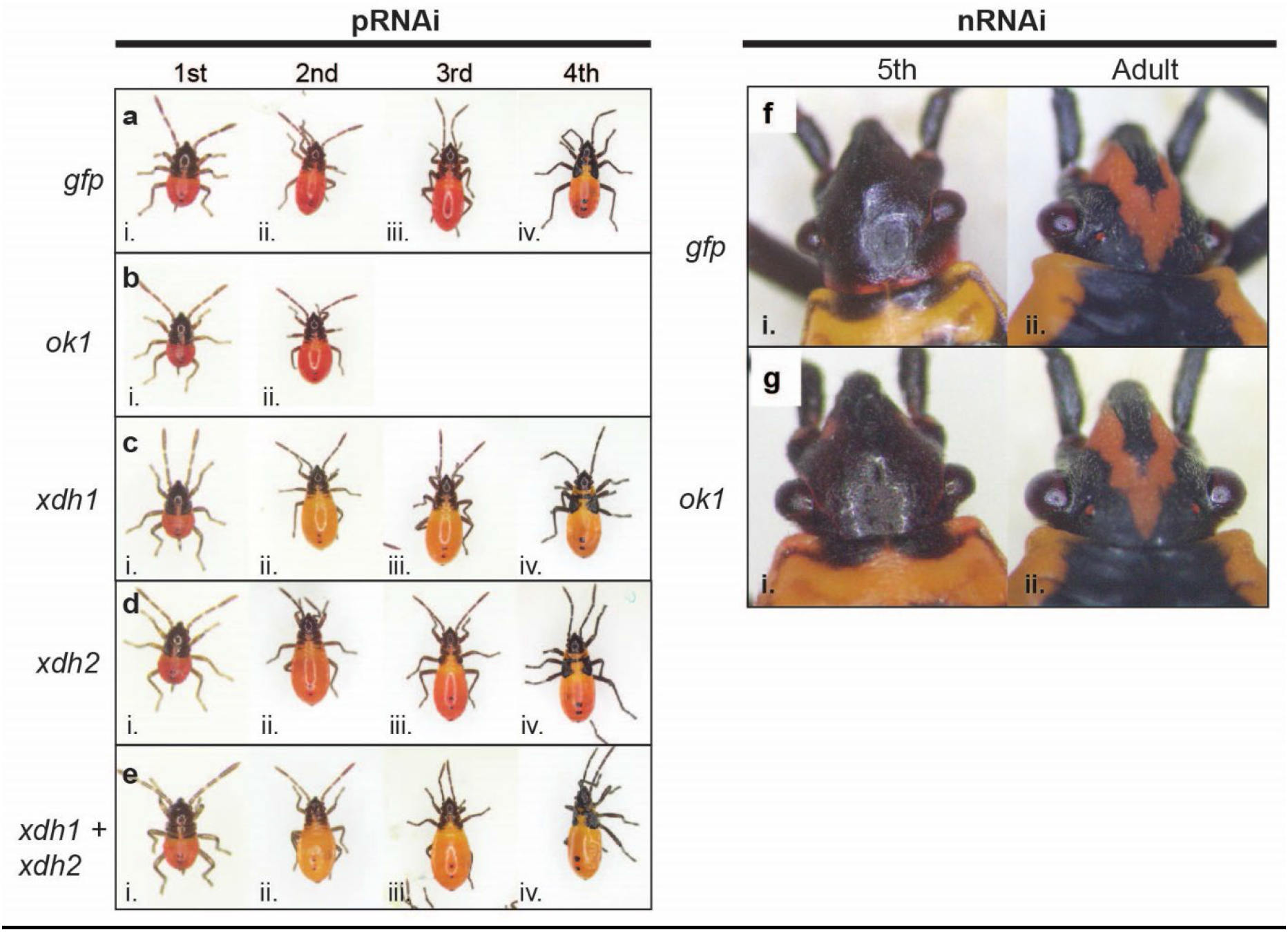
RNAi targeting pteridine synthesis pathway in *Oncopeltus*. Photos of nymphs and adults, as indicated. **(A)** Offspring of females injected with a *gfp* dsRNA control exhibit wild type phenotypes at (i) first, (ii) second, (iii) third, and (iv) fourth instars. (B) Offspring of females injected with *Of-ok1* dsRNA were indistinguishable from controls. pRNAi knockdown of (C) *Of-xdh1*, (D) *Of-xdh 2*, or (E) *Of-xdh 1* and *Of-xdh 2*. Knockdown of *Of-xdh 1* caused a lightening of body color from red (A) to yellow-orange, particular evident in second through fourth instars (Cii-iv; Eii-iv). (F-G) Nymphal RNAi. The eye color of last instar nymphs and adults are similar in controls (Fi, ii) and after nRNAi*^ok1^* (Gi, ii).

### xdh1 *contributes to body pigmentation in* O. fasciatus

In *Drosophila, rosy* functions downstream of *brown* in the pteridine pathway, and mutants display dark red eyes^26^. To further investigate potential components of the pteridine pathway in *O. fasciatus*, we used pRNAi to assess the functions of the *xanthine dehydrogenase (xdh)* ortholog to *Drosophila rosy*. BLAST searches of the *O. fasciatus* official gene set (OGS) followed by phylogenetic analysis (Fig. S1) revealed two orthologs of *Dmel-ry,* which we refer to as *Of-xdh1* (OFAS027123) and *Of-xdh2* (OFAS013815) (Fig. S1, S2). However, as we were unable to amplify the region encompassing both dsRNA templates of *Of-xdh2,* it is possible the OFAS013815 gene model encodes more than one transcript (see Methods for more detail). Knockdown of *Of-xdh1* by pRNAi produced nymphs with slightly lighter body coloration than *gfp*^pRNAi^ nymphs, appearing yellow-orange rather than red (Fig. 1A,C). This difference was most discernible by the 2^nd^ instar (Fig. 1Cii), likely because the compaction of the body in newly hatched 1^st^ instar nymphs obscures slight differences in pigmentation (Fig. 1Ci). The body color change persisted through the third and fourth nymphal instars (Fig. 1Ciii-iv). However, following *Of-xdh2* pRNAi, no clear change in pigmentation relative to *gfp* controls was observed at any instar, although a marginal decrease in red body color intensity cannot be ruled out (Fig. 1D). Simultaneous knockdown of both *Of-xdh1* and *Of-xdh2* resulted in nymphs with yellow-orange coloration indistinguishable from the *Of-xdh1* single knockdown (Fig. 1E). The persistence of the yellow-orange phenotype observed after *Of-xdh1* pRNAi through several molts, from 1^st^ to 4^th^ instar nymphs (Fig. 1C), indicates that pteridine pigments are unaffected by ecdysis since they are deposited in the epidermis as opposed to the cuticle^51,52^. However, some *Of-xdh1* dsRNA-treated nymphs appeared wild type by the fourth instar, indicating that the RNAi effect waned over time in some individuals.

Since we have noticed variation in body coloration within our *O. fasciatus* lab population and wanted to ensure that the lighter body coloration observed after *xdh1* pRNAi was indeed due to specific knockdown of this gene and not lighter variants already present in the population, we performed another round of *Of-xdh1* pRNAi to be scored blindly. In this experiment, adults were injected with either *gfp* or *Of-xdh1* dsRNA by one researcher, and second instar offspring from both treatments were individually placed in tubes labeled with a random number to be analyzed by a second blinded researcher, described in more detail in the Methods. The blinded researcher scored each individual using a four-point scale (yellow-orange, orange, orange-red, or red). Of 188 *gfp* pRNAi offspring scored, 97.8% were scored as orange-red or red; just 4 (2%) were scored as orange. Of 193 scored *Of-xdh1* pRNAi bugs scored, none were scored as red, just one (0.5%) was scored as orange-red, and 99.4% were scored as yellow-orange or orange. Thus, *Of-xdh1* knockdown produced a definitive and consistent reduction in body pigmentation outside the range of variation in seen in wild type populations. Overall, these results demonstrate that *Of-xdh1* is required for dark red pigments in the *O. fasciatus* body, consistent with a role in the pteridine synthesis pathway, and does not appear to be required for viability, making this a potentially useful marker in this species. Interestingly, we did not observe any effect of *Of-xdh1* on eye color.

### Of-v *and* Of-st *are required for brown pigmentation in adult compound eyes*

We next investigated roles of genes expected to function in ommochrome pigmentation pathways. The tryptophan 2,3-dioxygenase-coding gene *vermilion* (*v*) and ABC transporter-encoding gene *scarlet* (*st*) are involved in ommochrome pigment synthesis and transport in *Drosophila* and other insects (see Introduction). Furthermore, previous work showed that RNAi targeting either *Of-v* or *Of-st* results in loss of dark brown pigmentation in the eyes^47,48^. We employed both pRNAi and nRNAi to assess the function of *v* and *st* in *O. fasciatus*. Following pRNAi targeting either *Of-v* or *Of-st*, no change in eye or body coloration was observed relative to *gfp* controls in 1^st^ or 2^nd^ instar nymphs (Figure 2A-C). However, nRNAi targeting *Of-v* or *Of-st* in 4^th^ instar nymphs resulted in a loss of brown pigments at the medial edge of the compound eyes in 5^th^ instar nymphs (Fig. 2Ei,Fi). The effect was even more pronounced after individuals molted to adulthood (Fig. 2Eii, Fii). This partial change in eye color is attributed to the gradual addition of newly formed ommatidia after each molt. No change in body coloration following knockdown of *Of-v* or *Of-st* was observed. In short, *Of-v* and *Of-st* both appear to play roles in ommochrome synthesis or transport, respectively, with loss of these transcripts resulting in clear phenotypic differences late in development. In addition, no impact on viability was observed after RNAi knockdown for either gene (Table 1), making these genes good candidates for use as eye color markers in this species.

**Figure 2.**
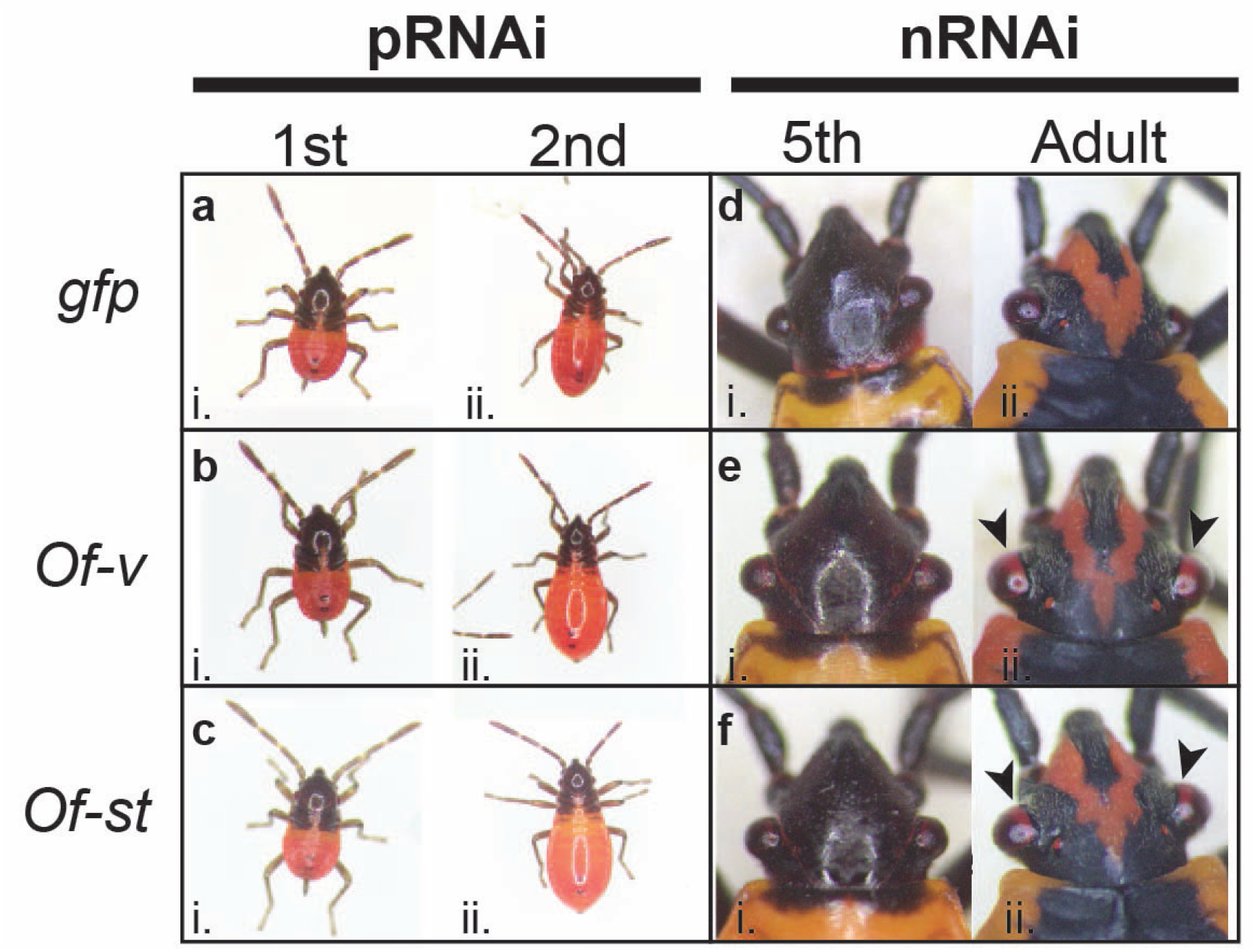
RNAi knockdown of ommochrome pathway genes *Of-st* and *Of-v* results in loss of brown eye pigmentation. Photos of nymphs and adults, as indicated. (A) Offspring of females injected with a *gfp* dsRNA control exhibit wild type phenotypes at first (i), second (ii) instars. Offspring of females injected with (B) *Of-v* dsRNA or (C) *Of-st* dsRNA. Fourth instar nymphs were injected with (D) *gfp*, (E) *Of-v*, or (F) *Of-st* dsRNA, shown after molting to fifth instar (i) and adulthood (ii). While *gfp* dsRNA-treated individuals display wild type black eyes, those treated with either *Of-v* or *Of-st* dsRNA display clear loss of brown eye pigment in the new ommatidia along the medial border of the adult compound eye (Eii, Fii, arrowheads).

### *Loss-of-function red-eyed* Of-vermilion *mutants are viable*

Given that RNAi knockdown of *Of-v* did not appear to negatively impact viability or fertility, we chose it as a candidate for generating a stable eye color marker for *O. fasciatus.* We used CRISPR/Cas9 to generate an *Of-v* null mutant, with tools we developed previously for mutagenesis in *O. fasciatus*^9^. We designed a single guide RNA (sgRNA) to direct Cas9 to a site within exon 2 of *Of-v* (Fig. 3A arrowhead; see Methods). Of 1177 embryos injected with *Of-v* gRNA and Cas9 protein, 779 (66%) hatched. Since it takes at minimum four months from the time of *O. fasciatus* embryo injection to generation of homozygous individuals when reared at 25°C, we sought to increase the developmental rate by rearing the G0s at 29°C and 75% relative humidity. At their final nymphal instar, we observed an unusually high number of dead G0 nymphs, all of which had swollen black abdomens, a presentation we had not previously seen in our lab populations (Fig. S3). While we cannot be sure of the cause, the symptoms we observed closely matched those described for a disease of *O. fasciatus* populations infected with the bacterium *Pseudomonas aeruginosa*^53^, namely: paralysis, darkening of the body, bad smell, and very quick death following the onset of symptoms. The disease caused by *P. aeruginosa* was likewise associated with populations raised at high temperature (30°C) and humidity.

**Figure 3.**
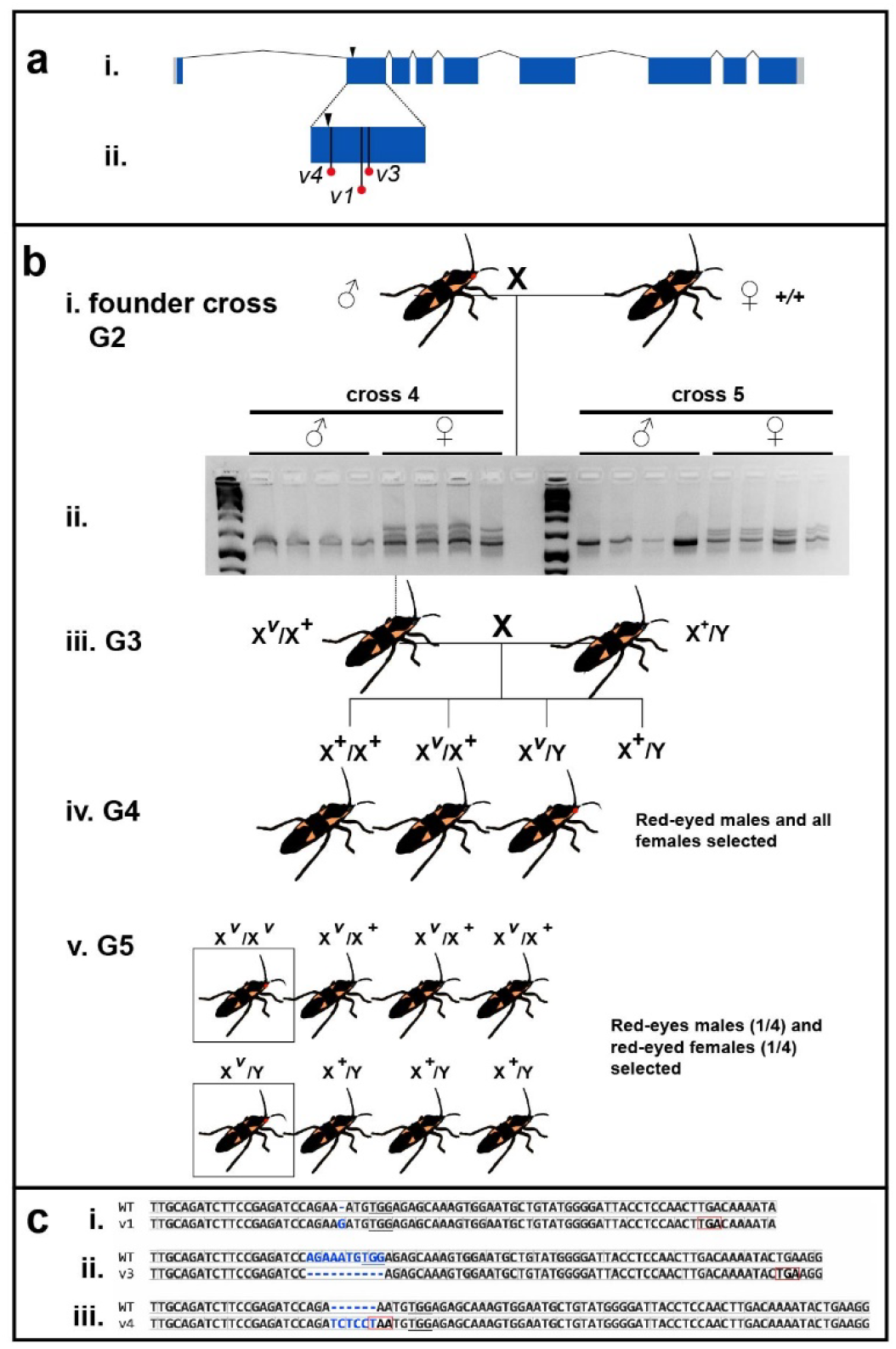
Generation of *Of-v* mutant lines using CRISPR/Cas9. (A) An *Of-v* gRNA was designed to guide Cas9 to generate a double-stranded break (arrowhead) in *Of-v* exon 2. (i) *Of-v* gene structure, with coding DNA sequence shown in dark blue and untranslated regions shown in light gray. (ii) Premature stop codons found in exon 2 of all mutant alleles. (B) Crossing scheme to generate homozygous lines from a heteroallelic *Of-v* mutant stock. (i) Five single crosses were set up between a single red-eyed *Of-v* G2 male founder (derived from the heteroallelic *Of-v* population) and three wild-type virgin females. (ii) *Of-v* exon 2 was amplified from genomic DNA isolated from the progeny of these crosses and a heteroduplex mobility assay performed; all females gave rise to heteroduplex bands, while no heteroduplex bands were found in male samples, consistent with *Of-v* X linkage; (iii) One G3 female offspring from each founder cross was mated with one or more wild type males; (iv) Male G4 offspring had either red or black eyes, while all female G4s had black eyes. All red-eyed males were selected to mate with female siblings; (v) In the G5, all red-eyed males and red-eyed virgin females were selected to establish a homozygous lines. The expected genotypes and phenotypes are shown in their expected proportions. (C) Alignment between the wild-type *Of-v* allele and the three mutant alleles for which homozygous lines were successfully established: *v^1^*, has a single nucleotide insertion at the predicted double-stranded break site (i), *v^3^*, which has a 10 bp deletion (ii), and *v^4^*, which has a 6 bp insertion. Insertions or deletions, blue. Reading frame, light gray boxes. Introduced stop codons, red boxes. PAM site, underline.

The eyes of healthy G0s, sequestered from the diseased population, were inspected for the expected red-eyed mosaic phenotype, which would result from biallelic hits in the injected animals. Mosaicism was observed in many individuals. In our efforts to isolate potentially infected bugs from the rest of our *O. fasciatus* lab population, we were unable to tabulate the frequencies of G0 phenotypes. Rather, all G0s with a red-eyed mosaic phenotype were allowed to mate to each other, producing many red-eyed G1s, which were selected and maintained as a heteroallelic *Of-v* mutant population.

### Of-vermilion *X-linkage and establishment of homozygous lines*

Having established a population of red-eyed mutant bugs with many different *v* mutant alleles, we sought to establish single-allele homozygous lines. Five G2 males, offspring of the red-eyed G1 heteroallelic *Of-v* stock described above, were individually outcrossed to wild type virgins (“founder crosses”, Fig. 3B). When founder offspring reached the 5^th^ instar or early adulthood, a single leg was removed for genomic DNA isolation, allowing us to genotype individuals before crossing them. We found that samples from all female offspring of crosses 3-5 showed the presence of a mutant allele, while no male samples produced heteroduplex bands, consistent with *Of-v* X-linkage (Fig. 3Bii). To test for X-linkage, a single red-eyed virgin female from the heteroallelic *Of-v* population was outcrossed to a wild-type male. When all progeny of this cross were in their final nymphal instar or adults, their eye color was examined. All female offspring (37/37) were found to have wild-type black eyes, and all male offspring (30/30) were found to have red eyes, consistent with X-linkage (data not shown).

Accordingly, to generate homozygous lines, a single virgin presumably heterozygous G3 female offspring from each of the original five founder crosses was outcrossed to one or more wild-type males (Fig. 3Biii). All of these crosses produced both red- and black-eyed G4 male progeny (Fig. 3Biv). Red-eyed males and all females from each cross were selected and crossed to each other. Finally, the G5 red-eyed progeny were selected to produce homozygous lines (Fig. 3Bv).

From the original five founder crosses, three homozygous *Of-v* mutant lines were established: *v^1^, v^3^,* and *v^4^*. Sequencing of *Of-v* exon 2 from each line showed all three alleles result in premature stop codons within exon 2 (Fig. 3Aii, red dots; Fig. 3C). The wild type *Of-v* allele has a 1167 bp long coding DNA sequence (CDS). Allele *v^1^* has a single guanine insertion at the expected double-stranded break (DSB) site, causing a frameshift and introduction of a stop codon at position 87 of the CDS (Fig. 3Ci). Allele *v^3^* has a 10 bp deletion, removing bases from both sides of the expected DSB, as well as introducing a stop codon at CDS position 98 (Fig. 3Cii). We also observed a single substitution in the intron preceding exon 2 in which a guanine was present where a thymine was observed in 8/8 wild-type samples. As this locus is 61 bp away from the expected DSB site, this difference most likely represents a polymorphism present in the G0 individual before Cas9 cleavage. Allele *v^4^* has a 6 bp (TC|TCC|T) insertion immediately before AA (Fig. 3Ciii), resulting in a TAA stop codon at that site (Fig. 3Ciii, red box).

### *Scoring* Of-v *mutants for use as genetic tool*

All three *Of-v* mutant lines are viable and fertile, making *Of-v* useful as an eye color marker for this species. To determine at which instars the *Of-v* mutant phenotype can reliably be scored, we photographed individuals from each of our three *Of-v* mutant lines at every nymphal instar and as adults to document and compare the phenotype to wild-type bugs over the lifespan. During the first instar (Fig. 4A), the *Of-v* lines are almost indistinguishable from wild type, though the body color of *Of-v* mutants is slightly lighter (Fig. 4Ai, dorsal view; ii, lateral view). Consistent with pRNAi results, the eye color of wild type first instar nymphs is not clearly different from that of the *Of-v* lines (Fig. 4Aiii-vi), perhaps because ommochromes have not yet been deposited in the eye. By the second instar, there is a clear difference in eye color between wild type (Fig. 4Biii) and *Of-v* mutant lines (Fig. 4Biv-vi). Any differences in body coloration become less apparent as development proceeds, while differences in eye pigmentation become much more pronounced as the compound eyes enlarge in subsequent instars (Fig. 4C-E). In adulthood, male and female *Of-v* mutants have bright red compound eyes in stark contrast to the black eyes of wild type adults (Fig. 4F, G). Overall, we noticed no differences in phenotype between the three *Of-v* strains. We have maintained each *Of-v* line for multiple generations and have noticed no difference in their developmental rate or overall health compared to our wild type population. Thus, these mutants provide a reliable visible marker for the *O. fasciatus* research community for transgenesis and other genetic manipulations.

**Figure 4.**
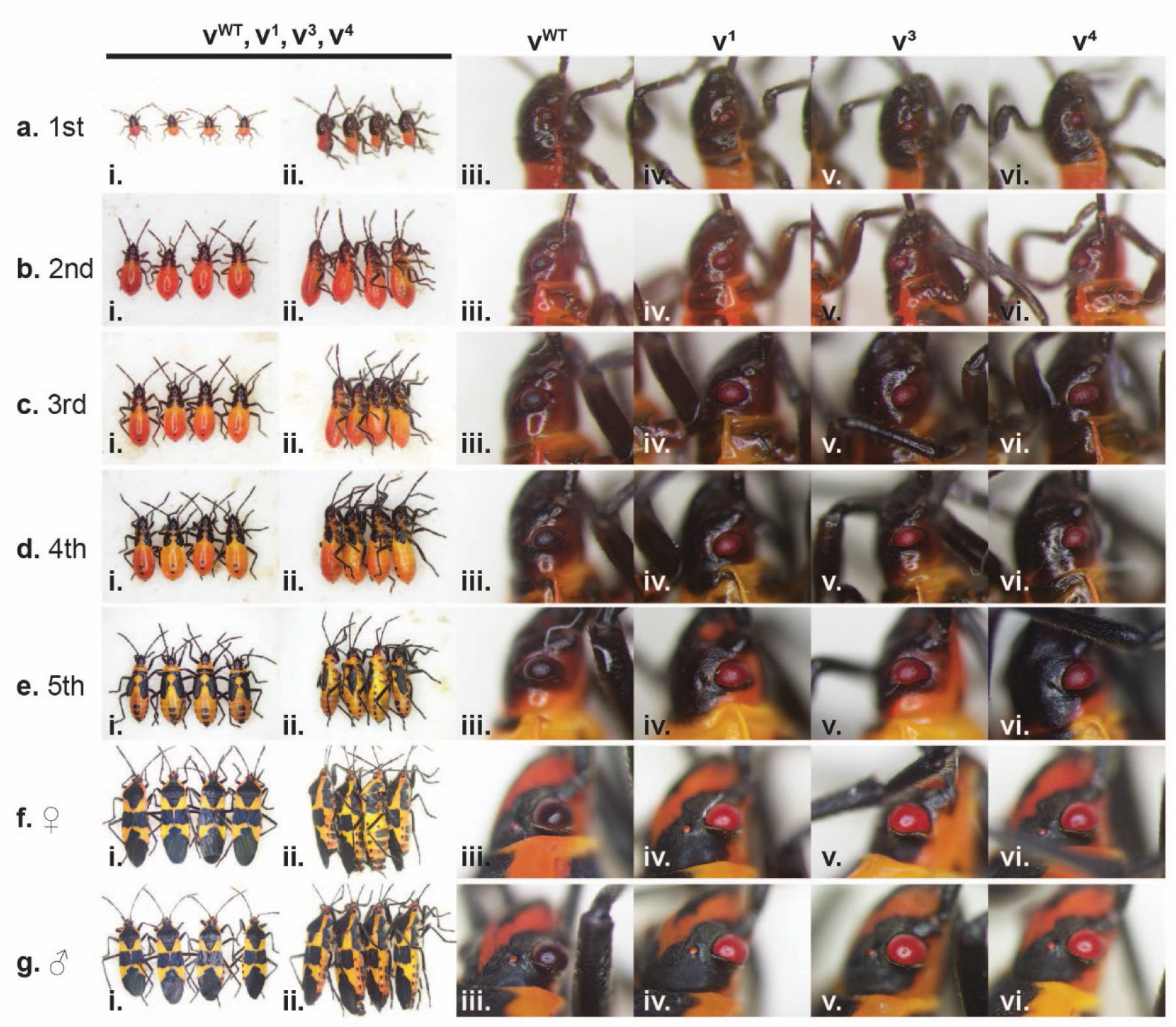
Phenotypes of homozygous *Of-v* lines across lifespan. Photographs of whole animals or heads of different individuals are shown in A-G. (A) First instar nymphs, (B) Second instar nymphs, (C) Third instar nymphs, (D) Fourth instar nymphs, (E) Fifth instar nymphs, (F) Adult females, (G) Adult males. (i.) Dorsal view of each genotype in the following order: v^WT^, v^1^, v^3^, v^4^. (ii.) Lateral view of each genotype in the same order as shown in i. (iii-vi.) Close-up view of right compound eye from individuals shown in i-ii, with genotypes indicated at top. Wild-type and *v* phenotypes are essentially indistinguishable during the first instar (A), but differences in coloration become apparent as wild-type eyes darken by the second instar (B), while *v* compound eyes remain red for the rest of nymphal development (C-E) and appear bright red in adults (F-G).

Because of issues with disease described above, we were not able to determine the frequency of mutagenesis in our initial *Of-v* CRISPR experiment. To determine basic statistics of CRISPR/Cas9 efficiency and germline mutation rates, we conducted a second round of *Of-v* CRISPR injections, making use of our new *Of-v* mutant lines for crosses (Fig. S4). Wild type embryos were injected with Cas9 protein and an *Of-v* gRNA as described above. We characterized the phenotype of mosaic G0s, and crossed these to *v^4^* virgins. G1 offspring from each cross were then analyzed for red or black eyes. We observed a progressive increase in germline mutation rates with the severity of somatic knockout seen in the mosaic G0s. G0s that appeared wild type produced a germline mutation rate of 66.18±9.12%, while G0s with one red eye had a germline mutation rate of 74.56±9.37%, and those with two red eyes had a rate of 90.41±4.33% (Table 2). Importantly, these results suggest that *Of-v* co-mutation may be a helpful strategy to concentrate screening efforts when trying to isolate non-visible mutations at other loci.

**Table 2.** Somatic and germline mutation rates observed in second *Of-v* CRISPR experiment.

| **G0 phenotype** | **Number** | **Percent**<br>**of all G0s** | **Mean**<br>**germline**<br>**mutation**<br>**rate** | **SEM**<br>**germline**<br>**mutation**<br>**rate** |
| --- | --- | --- | --- | --- |
| WT (total) | 21 | 30.43 | 66.18 | 9.12 |
| WT (males) | 9 | 13.04 | 45.93 | 16.04 |
| WT (females) | 12 | 17.39 | 81.38 | 8.67 |
| 1 red eye (total) | 14 | 20.29 | 74.56 | 9.37 |
| 1 red eye (males) | 5 | 7.25 | 63.90 | 20.82 |
| 1 red eye (females) | 9 | 13.04 | 89.81 | 4.96 |
| 2 red eyes (total) | 34 | 49.28 | 90.41 | 4.33 |
| 2 red eyes (males) | 24 | 34.78 | 89.09 | 5.69 |
| 2 red eyes (females) | 10 | 14.49 | 93.56 | 5.80 |

### xdh1 *knockdown in v^3^ mutant produces bright yellow body color*

Given that we observed a lighter body color after *Of-xdh1* pRNAi (Fig. 1C) and after *Of-v* mutation (Fig. 4A-C), we wondered if both genes might contribute to body pigmentation in an additive manner. To test this, we performed *Of-xdh1* pRNAi in an *Of-v* mutant background. *Of*-*v^3^* females were injected with either *Of-xdh1* dsRNA or *gfp* dsRNA as a negative control. Interestingly, *v^3^; Of-xdh1* pRNAi offspring had bright yellow bodies (Fig. 5B), lighter than the *v^3^; gfp*^pRNAi^ controls (Fig. 5A), and even lighter than the yellow-orange pigmentation observed after *xdh1* pRNAi knockdown in a wild type background (compare to Fig. 1C). No distinguishable effect of the knockdown on *Of*-*v^3^* eye pigmentation was observed, consistent with our previous observation that *Of-xdh1* knockdown in a wildtype background had no effect on eye pigmentation. The yellow body phenotype was marginally detectable in 1^st^ instar nymphs (Fig. 5Bi) but was clear in 2^nd^ instars (Fig. 5Bii) and lasted until through the 4th instar (Fig. 5Biv). These results suggest that both *Of-v* and *Of-xdh1* contribute independently to *O. fasciatus* body coloration and that *Of-xdh1* would be useful as a visible body color marker for *O. fasciatus*.

**Figure 5.**
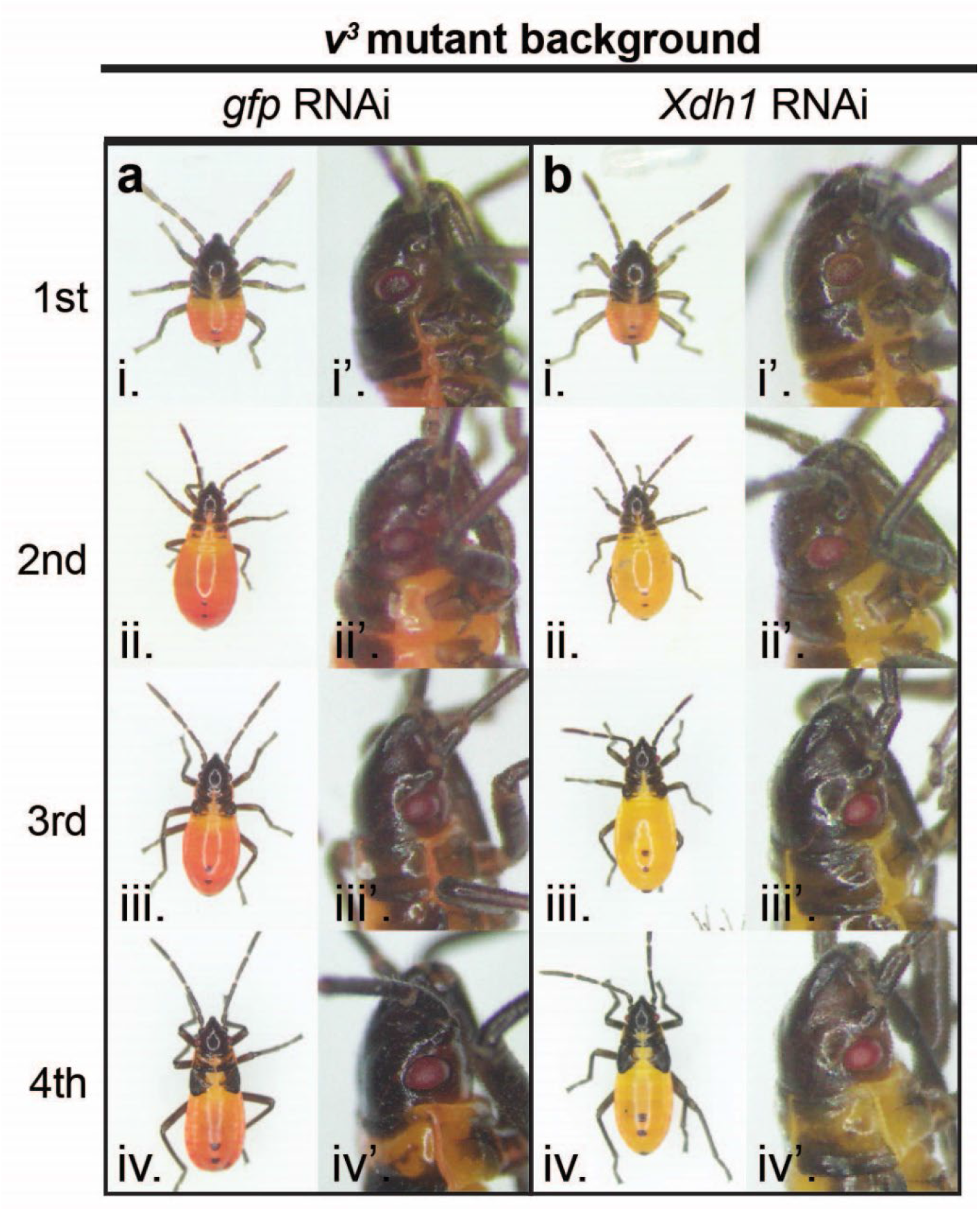
*Of-xdh1* knockdown in *Of-v3* mutant background produces bright yellow body color. Offspring of v^3^ mothers treated with *gfp* (A) or *Of-xdh1* (B) dsRNA were observed at first (i), second (ii), third (iii), and fourth (iv) instars. While the knockdown did not appear to produce an effect in the eyes of pRNAi offspring (i’-iv’), body pigmentation was greatly reduced in *Of-xdh1* treated individuals, resulting in a bright yellow color (Bi-iv).

## Discussion

Here we have expanded the repertoire of genetic tools available for *O. fasciatus*, an emerging model system for hemimetabolous insects. Among pigmentation mutants, those that display loss of eye pigmentation are particularly valuable, given the established efficacy the 3XP3 synthetic enhancer-promoter construct, which can direct gene expression in eyes, across a range of phylogenetically diverse insects^54^. Combined with a fluorescent protein-encoding gene or rescue allele, this becomes a powerful tool for screening low-frequency genetic events, including gene replacement using CRISPR/Cas9 or transposon-mediated transgenesis. Even in the absence of validated transgenes, pigmentation mutants are useful for tracking germline mutation in the case of CRISPR/Cas9 co-conversion, allowing alleles that may not produce visible phenotypes to be more easily detected.

In investigating genes that would be useful as visible markers, we identified functions for both ommochrome and pteridine synthesis pathways in eye and body color of these bugs. This contrasts with holometabolous insects such as the beetle *Tribolium castaneum*, where mutations in ommochrome pathway genes generate white-eyed animals, suggesting no role for pteridines in eye color in that species^55,56^. Based on our previous experience that loss-of-function mutation of *Of-w* results in lethality to homozygotes^9^, we pre-screened candidate genes by RNAi before performing CRISPR/Cas9 mutagenesis. Similar to previous work in *O. fasciatus*^47^ and *R. prolixus*^40^, we did not observe roles for *Of-ok* paralogs in eye pigmentation, but also observed no change in body pigmentation following *Of-ok1* knockdown, in contrast to the functions of these genes in *R. prolixus*. However, we did observe an unexpected role for *Of-ok2* in embryonic viability. Knockdown of *Of-xdh1*, an ortholog of *Drosophila rosy*, revealed a subtle but consistent change in body color, with loss of red pigmentation, while no role was identified for *Of-xdh2* (Fig. 1). Within the ommochrome pathway, our studies confirmed roles of *Of-st*, encoding a putative ommochrome transporter, and *Of-v*, encoding tryptophan 2,3-dioxygenase, in eye color (Fig. 2). Neither parental nor nymphal RNAi appeared to impact viability, making both of these genes potentially useful eye color markers.

Accordingly, we used CRISPR/Cas9 genome editing to generate three independent loss-of-function mutations via non-homologous end joining in *Of-v* (Fig. 3,4), each of which has been maintained as a viable colony of bright red-eyed homozygotes for multiple generations without any apparent impact on viability or fertility. As in *Drosophila*, this gene is located on the X-chromosome of *O. fasciatus* (Fig. 3). Thus, although the phenotype appears identical to that of Lawrence’s *re* mutants, they are not allelic as *re* behaved as an autosomal recessive^44^. We further made use of *Of-v* mutants to test the efficiency of CRISPR/Cas9 mutagenesis in this species, showing that bugs exhibiting mosaicism in G0 individuals had a 90% rate of germline transmission, making *Of-v* an excellent marker for co-CRISPR^57^. Finally, using RNAi to knockdown *Of-xdh1* in *Of-v* homozygotes, we revealed a role for the ommochrome pathway in body coloration, as the body color of these double “mutants” was bright yellow (Fig. 5), compared to the yellow-orange color of *Of-xhd1*^RNAi^ animals (Fig. 1). *Of-xdh1* does not appear to be required for viability, making this another potentially useful *O. fasciatus* marker. In contrast to our findings, studies in water striders suggested that ommochromes play no role in body pigmentation but function only for eye color^41^. Conversely, *rosy* plays roles in body and eye pigmentation in water striders^41^, but we found it to function only in body coloration in *O. fasciatus*. These differences may reflect species variation or stage-specific utilization of the different pathways, which remain to be examined.

## Methods

### O. fasciatus *rearing*

An *O. fasciatus* wild type population was maintained at room temperature (23-24°C) in clear plastic storage containers (15.25×11×11 inch) and exposed to a 16h:8h light:dark cycle. All RNAi experiments were performed at 25°C. As described in the Results, subpopulations were briefly maintained at 29°C and 75% relative humidity, which appeared to promote disease (Fig. S3). A paper towel and cotton ball wick placed atop a 32 oz. plastic container full of water provided a constant source of water and raw organic sunflower seeds were provided as food. Single crosses were performed either using *Drosophila* vials as described^9^ or using one gallon plastic storage containers.

### Gene isolation

To identify the *O. fasciatus* ortholog of *v,* a BLAST search against the *O. fasciatus* official gene set (OGS)^8^ was conducted using the *Dmel-v* amino acid sequence as a query. Only one hit had an e-value less than 1.4, that of OFAS025214 (e-value = 2e-129), suggesting that there is only one *Of-v* ortholog. To confirm the identity of OFAS025214, a phylogenetic tree was constructed from an alignment of this gene model to amino acid sequences of several insect *v* orthologs. The OFAS025214 gene model clustered with the *Lhes-v,* the only other heteropteran *v* ortholog included in this analysis, suggesting that OFAS025214 is indeed the *Of-v* ortholog. To identify the *Of-xdh* ortholog, the *O. fasciatus* OGS was input into BLAST using the *Dmel-ry* amino acid sequence as query. Several hits had significant e-values, so sequences for all gene models with e-values below 8e-71 were collected and subjected to phylogenetic analysis. The resulting gene tree showed two gene models—OFAS027123 and OFAS013815—clustering with other insect *xdh/ry* orthologs, whereas the remaining *O. fasciatus* gene models clustered with *Dmel-aox,* which was included in the analysis an outgroup, suggesting that there are two *Of-xdh* paralogs. The *O. fasciatus* orthologs of *st, ok1,* and *ok2* were analyzed previously^9^. Primers to all sequences were designed based on *O. fasciatus* genome sequence (see Table S1). For *Of-v,* we amplified the coding DNA sequence using primers 30 and 31, and the PCR product was blunt-end ligated into SmaI-cut pUC19, and subsequently Sanger sequenced. We Sanger sequenced fragments from all other genes spanning the two dsRNA templates (Fig. S2), except for *Of-xdh2.* For this gene model (OFAS013815), we were able to amplify two non-overlapping fragments, but we were not able to amplify a fragment uniting these two regions. This may suggest that the two regions are not part of the same transcript; more work is needed to determine if the OFAS013815 gene model represents a single gene.

### Phylogenetic analysis

To construct gene trees, amino acid sequences were aligned in MUSCLE^58^ followed by phylogenetic inference using MrBayes 3.1.1^59^ within the TOPALi v2.5 framework^60^ with the following parameters: runs = 2, generations = 100000, sample freq. = 10, burnin= 25%. Tree formatting was done in MEGA X 10.1.6^61^.

### Double-stranded RNA (dsRNA) synthesis

All dsRNA templates were PCR-amplified using Q5 high-fidelity DNA polymerase (NEB) from *O. fasciatus* first instar complementary DNA (cDNA). In some cases, this PCR product was inserted into the pGEM T-easy vector before dsRNA synthesis, while in others the PCR product was used directly as template for dsRNA synthesis. A dsRNA template targeting *turbo gfp (tGFP)* was amplified from plasmid pBac[3xP3-DsRed; UAS-Tc’hsp_p-tGFP-SV40] (Addgene 86453) for use as a negative control. All primers used to amplify dsRNA templates have T7 RNA polymerase promoter sequences at each 5’ end allowing synthesis of both RNA strands in a single reaction. PCR products were used as templates in an RNA transcription reaction using the MEGAscript T7 Transcription kit (Invitrogen) at 37°C overnight. 1 ul kit-provided TURBO DNase was then added and the reaction was incubated at 37°C for 15 min. The RNA was denatured by heating to 95°C for 3 min in a heat block; the heat block was then turned off, and the RNA strands were allowed to anneal as the heat block temperature slowly decreased. The dsRNA was precipitated with ethanol and lithium chloride, and resuspended in water. For each gene except *Of-v* and *tGFP,* two non-overlapping dsRNAs were made (denoted dsRNA A and B, Fig. S2). All primer sequences are listed in Table S1.

### Parental RNA interference (pRNAi)

For pRNAi, each dsRNA was diluted to 5 uM in injection buffer (5 mM KCl, 0.1 mM phosphate buffer pH 6.8) with green food coloring (McCormick) diluted 1:50. Each adult female was injected with 3 ul injection mix between the 4^th^ and 5^th^ abdominal sternites using a glass needle pulled from borosilicate glass capillary tubes (World Precision Instruments). Each injected female was paired with a male and kept individually in a *Drosophila* vial supplied with raw sunflower seeds, damp cotton as a source of water, and dry cotton for egg laying. Vials were laid on their sides to keep seeds from contacting the wet cotton. Embryos were collected daily over 7 days after injection and incubated at 25°C for approximately one week until hatching. 1^st^ and 2^nd^ instar nymphs of each group were photographed. For *Of-xdh1+xdh2* and *Of-ok1+ok2* double injections, either 2.5 uM of each dsRNA (dsRNA A, Figure S2) or 5 uM of each dsRNA (dsRNA B, Figure S2) were used.

After an initial *xdh1* knockdown experiment produced a subtle phenotype, we carried out a blind experiment to ensure that this phenotype was indeed due to *xdh1* knockdown and not to lighter colored variants in our population. In this experiment, phenotypes of offspring of *Of-xdh1* or *gfp* dsRNA-injected females were scored by a researcher who had no knowledge of the treatment each bug had received. Two researchers independently injected five adult females each with *Of-xdh1*-B dsRNA or *gfp* dsRNA (as described above). The first four egg clutches laid by each female were discarded as we have often observed that the first several clutches fail to display knockdown phenotypes, likely due to their position within the ovariole at the time of injection. Each additional clutch of embryos was placed in a *Drosophila* vial provided with sunflower seeds and damp cotton and incubated at 25°C for approximately 7 days until hatching. Nymphs were allowed to molt into 2^nd^ instar as we had noticed the lighter coloration seen after *xdh1* knockdown is more apparent at this stage. Each researcher selected 100 *gfp* pRNAi and 100 *Of-xdh1* pRNAi nymphs derived from their independent injections and placed each nymph in a tube labeled with a random number from 1 to 200. Each researcher gave the other researcher (the “analyzer”) the 200 nymphs to score on a 4-point scale from light orange to dark red. Any individuals found by the analyzer to be dead or at the wrong instar were excluded from analysis. The analyzer had no information about any of the nymphs aside from their random labels. Finally, the score sheets of the two analyzers were combined, treatment groups were unblinded, and scores assigned to individuals from each RNAi treatment group were tabulated and compared.

### Nymphal RNA interference (nRNAi)

For nRNAi, 4^th^ instar nymph injections were performed as described above for pRNAi, except that only 1 ul of the 5 uM dsRNA injection mix was injected into each nymph and individuals were injected between the 5^th^ and 6^th^ abdominal sternites. After injection, nymphs were kept in small cages provided with damp cotton and sunflower seeds, incubated at 25°C, and were photographed after molting to 5^th^ instar and/or adulthood.

### *CRISPR/Cas9 mutagenesis of* Of-vermilion

The *Of-v* exon 2 sequence was submitted to CHOPCHOP^62^ to identify potential gRNA target sites. Each of the suggested gRNAs was aligned to the *O. fasciatus* genome assembly using blastn -task blastn-short, and the resulting alignments were used to evaluate the potential of off-target Cas9 cleavage. A gRNA was selected that produced no alignments in the PAM plus seed region for any genomic region other than *Of-v* exon 2. This gRNA was synthesized in vitro following the recommendations of https://www.crisprflydesign.org/grnatranscription/. Briefly, the gRNA sequence was introduced using primer 24, which was used with primer 23 to amplify a gRNA scaffold sequence in plasmid pCFD3^62^ using Phusion polymerase. This PCR product was added as template to a T7 RNA polymerase transcription reaction using the Megascript T7 kit (Invitrogen). Following 3 h of RNA transcription at 37°C, TURBO DNase was added to the reaction and incubated for 15 min at 37°C. The RNA was purified using the Monarch RNA Cleanup kit (NEB). To form ribonucleoprotein complexes prior to injection, the *Of-v* gRNA was diluted in injection buffer (5 mM KCl, 0.1 mM phosphate buffer pH 6.8) and heated to 70°C for 2 min, then transferred to ice. Cas9-NLS protein (PNA Bio) was added to the gRNA and incubated at room temperature for 5 min. The final injection mix contained 80 ng/ul *Of-v* gRNA, 300 ng/ul Cas9 protein, and green food coloring (McCormick) diluted 1:50. *O. fasciatus* embryos were collected over a ~3.5 h period and were injected at ~3-7 h AEL. Embryo injection was performed as described in Reding and Pick (2020).

### Heteroduplex mobility assay

When sampling 5^th^ instar or adult bugs, genomic DNA was prepared as described in ^63^, except that a single mesothoracic leg was used instead of whole individuals and only 25 ul of squishing buffer was used, which was then diluted 1:2 with water after DNA extraction. When sampling embryos, a single embryo was squished in 6 ul squishing buffer. Primers 25 and 26 were used to amplify a 581 bp fragment around *Of-v* exon 2, and the heteroduplex mobility assay was performed following standard methods ^9,64^.

### *Crosses to establish homozygous* Of-v *lines*

All G0s were inspected as young adults or last instar nymphs to identify mosaic animals, reflecting biallelic gene targeting events. As described in the Results, G0s with at least one red eye were selected and allowed to mate with each other. All G1s with two red eyes – presumably homozygous for *Of-v* mutation – were selected to establish a heteroallelic *Of-v* mutant population. Five G2 red-eyed males drawn from this population were each crossed to three wild-type virgins (Fig. 4i). A heteroduplex mobility assay was performed on embryos from the offspring of these crosses. Since this assay is not sensitive enough to detect very small indels, it was expected that not all samples would give rise to heteroduplex bands; indeed no heteroduplex bands were observed in samples of offspring from two of the five crosses. Offspring from three of the five founder crosses produced heteroduplex bands in all female fifth instar or adult samples and in none of the male samples, suggesting that *Of-v* is X-linked. Crosses proceeded as described in the Results and illustrated in Fig. 3B. To characterize each line’s mutation, genomic DNA was isolated as described above from two individuals per line, *Of-v* exon 2 was independently PCR-amplified from each sample using primers 25 and 26, and PCR products were Sanger sequenced.

### *Second* Of-v *CRISPR experiment*

Wild type embryos were injected with 300 ng/ul Cas9 protein (PNA Bio), 500 ng/ul pBac[3xP3-EGFP;Tc’hsp5’-Gal4Delta-3’UTR] (Addgene #86449), 80 ng/ul *Of-v* gRNA described above (see “CRISPR/Cas9 mutagenesis of *Of-vermilion*”), and 80 ng/ul of a gRNA designed to target the ampicillin resistance gene of the injected plasmid for Cas9-mediated linearization. This *AmpR* gRNA was synthesized in vitro as described above for the *Of-v* gRNA, except that primers 23 and 29 (Table S1) were used to amplify the gRNA template. The *AmpR* gRNA was injected to allow whole plasmid integration through non-homologous end joining at the *Of-v* locus, potentially tagging an *Of-v* mutation with a 3xP3>*egfp* fluorescent eye marker.

The phenotypes of adult G0s were recorded, before being crossed to *v^4^* virgins. 99 G0 crosses were set up; 30 G0s either died before mating or were infertile. G1s were CO2-anaesthetized and the eye phenotype of each was characterized as wild-type or *v* under a brightfield microscope; the presence of GFP in the eyes was assessed using a fluorescent microscope equipped with a GFP filter. Unfortunately, we observed no GFP^+^ G1 offspring, suggesting that either no plasmid incorporation occurred or that the reduction in eye pigmentation in *Of-v* mutants is not sufficient to allow GFP visualization. Germline mutation rate was differentially calculated for G0 males and G0 females, since male *v* G1 offspring of male G0s obtained their mutant *v* allele from their *v* mothers, and thus are not informative of *v* mutations that may have occurred in the paternal germline. Germline mutation rate for male G0s was calculated as the number of *v* female G1s / (total G1s – *v* male G1s); for G0 females the calculation was *v* G1s / total G1s.

## Acknowledgements

This work was supported by the National Institutes of Health (grant R01GM113230 to L.P.). K.R. was supported by a National Sciences Foundation Graduate Research Program Fellowship. pBac[3xP3-DsRed; UAS-Tc’hsp_p-tGFP-SV40] (Addgene 86453) and pBac[3xP3-EGFP;Tc’hsp5’-Gal4Delta-3’UTR] (Addgene 86449) were gifts from Gregor Bucher. We thank Patricia Graham and Matthew D. Fischer for comments on the manuscript.

## Author Contributions

K.R. and M.L. carried out experiments. K.R., M.L. and L.P. analyzed the data and wrote the manuscript. L.P. obtained funding.

## Data Availability Statement

Mutant strains and materials are available upon request. All sequences will be submitted to NCBI upon manuscript acceptance.

## References

1. Stork, N. E. How Many Species of Insects and Other Terrestrial Arthropods Are There on Earth? Annu Rev Entomol 63, 31–45, doi:10.1146/annurev-ento-020117-043348 (2018).

2. Panfilio, K. A. & Angelini, D. R. By land, air, and sea: hemipteran diversity through the genomic lens. Curr Opin Insect Sci 25, 106–115, doi:10.1016/j.cois.2017.12.005 (2018).

3. Jockusch, E. L. & Fisher, C. R. Something old, something new, something borrowed, something red: the origin of ecologically relevant novelties in Hemiptera. Curr Opin Genet Dev 69, 154–162, doi:10.1016/j.gde.2021.04.003 (2021).

4. Nunes-da-Fonseca, R., Berni, M., Tobias-Santos, V., Pane, A. & Araujo, H. M. Rhodnius prolixus: From classical physiology to modern developmental biology. Genesis 55, doi:10.1002/dvg.22995 (2017).

5. Pacheco, I. D., Walling, L. L. & Atkinson, P. W. Gene Editing and Genetic Control of Hemipteran Pests: Progress, Challenges and Perspectives. Front Bioeng Biotechnol 10, 900785, doi:10.3389/fbioe.2022.900785 (2022).

6. Hughes, C. L. & Kaufman, T. C. RNAi analysis of Deformed, proboscipedia and Sex combs reduced in the milkweed bug Oncopeltus fasciatus: novel roles for Hox genes in the hemipteran head. Development 127, 3683–3694, doi:10.1242/dev.127.17.3683 (2000).

7. Chipman, A. D. Oncopeltus fasciatus as an evo-devo research organism. Genesis 55, doi:10.1002/dvg.23020 (2017).

8. Panfilio, K. A. et al. Molecular evolutionary trends and feeding ecology diversification in the Hemiptera, anchored by the milkweed bug genome. Genome Biol 20, 64, doi:10.1186/s13059-019-1660-0 (2019).

9. Reding, K. & Pick, L. High-Efficiency CRISPR/Cas9 Mutagenesis of the white Gene in the Milkweed Bug Oncopeltus fasciatus. Genetics 215, 1027–1037, doi:10.1534/genetics.120.303269 (2020).

10. Morgan, T. H. Sex Limited Inheritance in Drosophila. Science 32, 120–122, doi:10.1126/science.32.812.120 (1910).

11. Sturtevant, A. H. The use of mosaics in the study of the developmental effect of genes. Proc. 6th Int. Congress of Genetics 1, 304–307 (1932).

12. Beadle, G. W. & Ephrussi, B. Development of Eye Colors in Drosophila: Transplantation Experiments with Suppressor of Vermilion. Proc Natl Acad Sci U S A 22, 536–540, doi:10.1073/pnas.22.9.536 (1936).

13. Beadle, G. W. & Ephrussi, B. The Differentiation of Eye Pigments in Drosophila as Studied by Transplantation. Genetics 21, 225–247, doi:10.1093/genetics/21.3.225 (1936).

14. Beadle, G. W. & Ephrussi, B. Development of Eye Colors in Drosophila: Diffusible Substances and Their Interrelations. Genetics 22, 76–86, doi:10.1093/genetics/22.1.76 (1937).

15. Ephrussi, B. & Beadle, G. W. Development of Eye Colors in Drosophila: Production and Release of cn Substance by the Eyes of Different Eye Color Mutants. Genetics 22, 479–483, doi:10.1093/genetics/22.5.479 (1937).

16. Ephrussi, B. & Beadle, G. W. Development of Eye Colors in Drosophila: Transplantation Experiments on the Interaction of Vermilion with Other Eye Colors. Genetics 22, 65–75, doi:10.1093/genetics/22.1.65 (1937).

17. Ephrussi, B. Chemistry of “eye color hormones” of *Drosophila*. The Quarterly Review of Biology 17, 327–338 (1942).

18. Popadic, A. & Tsitlakidou, D. Regional patterning and regulation of melanin pigmentation in insects. Curr Opin Genet Dev 69, 163–170, doi:10.1016/j.gde.2021.05.004 (2021).

19. Ziegler, I. & Harmsen, R. The biology of pteridines in insects. Advances in Insect Physiology 6, 139–203 (1970).

20. Kikkawa, H. Mechanism of Pigment Formation in Bombyx and Drosophila. Genetics 26, 587–607, doi:10.1093/genetics/26.6.587 (1941).

21. Summers, K. M., Howells, A. J. & Pyliotis, N. A. Biology of eye pigmentation in insects. Advances in Insect Physiology 16, 119–166 (1982).

22. Sullivan, D. T., Kitos, R. J. & Sullivan, M. C. Developmental and genetic studies on kynurenine hydroxylase from Drosophila melanogaster. Genetics 75, 651–661, doi:10.1093/genetics/75.4.651 (1973).

23. Ryall, R. L. & Howells, A. J. Ommochrome biolsynthetic pathway of Drosophila melanpgaster: variations in levels of enzume activities and intermediates during adult development. Insect Biochem. 4, 47–61 (1974).

24. Howells, A. J., Summers, K. M. & Ryall, R. L. Developmental patterns of 3-hydroxykynurenine accumulation in white and various other eye color mutants of Drosophila melanogaster. Biochem Genet 15, 1049–1059, doi:10.1007/BF00484496 (1977).

25. Figon, F. & Casas, J. Ommochromes in invertebrates: biochemistry and cell biology. Biol Rev 94, 156–183, doi:10.1111/brv.12441 (2019).

26. Reaume, A. G., Knecht, D. A. & Chovnick, A. The rosy locus in Drosophila melanogaster: xanthine dehydrogenase and eye pigments. Genetics 129, 1099–1109, doi:10.1093/genetics/129.4.1099 (1991).

27. Kim, H., Kim, K. & Yim, J. Biosynthesis of drosopterins, the red eye pigments of Drosophila melanogaster. IUBMB Life 65, 334–340, doi:10.1002/iub.1145 (2013).

28. Ewart, G. D. & Howells, A. J. ABC transporters involved in transport of eye pigment precursors in Drosophila melanogaster. Methods Enzymol 292, 213–224, doi:10.1016/s0076-6879(98)92017-1 (1998).

29. Mount, S. M. Sequence similarity. Nature 325, 487, doi:10.1038/325487c0 (1987).

30. Dreesen, T. D., Johnson, D. H. & Henikoff, S. The brown protein of Drosophila melanogaster is similar to the white protein and to components of active transport complexes. Mol Cell Biol 8, 5206–5215, doi:10.1128/mcb.8.12.5206-5215.1988 (1988).

31. Tearle, R. G., Belote, J. M., McKeown, M., Baker, B. S. & Howells, A. J. Cloning and characterization of the scarlet gene of Drosophila melanogaster. Genetics 122, 595–606, doi:10.1093/genetics/122.3.595 (1989).

32. Pepling, M. & Mount, S. M. Sequence of a cDNA from the Drosophila melanogaster white gene. Nucleic Acids Res 18, 1633, doi:10.1093/nar/18.6.1633 (1990).

33. Mackenzie, S. M. et al. Mutations in the white gene of Drosophila melanogaster affecting ABC transporters that determine eye colouration. Biochim Biophys Acta 1419, 173–185, doi:10.1016/s0005-2736(99)00064-4 (1999).

34. Dermauw, W. & Van Leeuwen, T. The ABC gene family in arthropods: comparative genomics and role in insecticide transport and resistance. Insect Biochem Mol Biol 45, 89–110, doi:10.1016/j.ibmb.2013.11.001 (2014).

35. Wang, L. et al. Mutation of a novel ABC transporter gene is responsible for the failure to incorporate uric acid in the epidermis of ok mutants of the silkworm, Bombyx mori. Insect Biochem Mol Biol 43, 562–571, doi:10.1016/j.ibmb.2013.03.011 (2013).

36. Liu, J., Lemonds, T. R., Marden, J. H. & Popadic, A. A Pathway Analysis of Melanin Patterning in a Hemimetabolous Insect. Genetics 203, 403–413, doi:10.1534/genetics.115.186684 (2016).

37. Brent, C. S. & Hull, J. J. RNA interference-mediated knockdown of eye coloration genes in the western tarnished plant bug (Lygus hesperus Knight). Arch Insect Biochem Physiol 100, e21527, doi:10.1002/arch.21527 (2019).

38. Heu, C. C. et al. CRISPR-mediated knockout of cardinal and cinnabar eye pigmentation genes in the western tarnished plant bug. Sci Rep 12, 4917, doi:10.1038/s41598-022-08908-4 (2022).

39. Xue, W. H. et al. CRISPR/Cas9-mediated knockout of two eye pigmentation genes in the brown planthopper, Nilaparvata lugens (Hemiptera: Delphacidae). Insect Biochem Mol Biol 93, 19–26, doi:10.1016/j.ibmb.2017.12.003 (2018).

40. Berni, M. et al. Atypical strategies for cuticle pigmentation in the blood-feeding hemipteran Rhodnius prolixus. Genetics 221, doi:10.1093/genetics/iyac064 (2022).

41. Vargas-Lowman, A. et al. Cooption of the pteridine biosynthesis pathway underlies the diversification of embryonic colors in water striders. Proc Natl Acad Sci U S A 116, 19046–19054, doi:10.1073/pnas.1908316116 (2019).

42. Abbott, C. E. Inherited melanism in the large milkweed bug, Oncopeltus fasciatus (Heteropters:Lygaeidae). Annals Entomological Society America 61, 542 (1968).

43. Smissman, E. E. & Orme, J. P. R. A yellow mutant strain of the large milkweed bug, *Oncopeltus fasciastus*, that lacks erythropterin. Annals Entomological Society America 62, 246 (1969).

44. Lawrence, P. A. Some New Mutants of Large Milkweed Bug Oncopeltus-Fasciatus Dall.Genet Res 15, 347–&, doi:Doi 10.1017/S0016672300001713 (1970).

45. Forrest, H. S., Menaker, M. & Alexander, J. Studies in the milkweed bug Oncopeltus fasciatus (Dallas). J. Insect Physiology 12, 1411–1421 (1966).

46. Shelton, P. M. J. & Lawrence, P. A. Structure and Development of Ommatidia in Oncopeltus-Fasciatus. J Embryol Exp Morph 32, 337–353 (1974).

47. Liu, J. Unraveling the molecular mechanisms of aposematic pigmentation in Oncopeltus fasciatus Ph.D. thesis, (2016).

48. Francescutti, C. M., Martin, A. & Hanly, J. J. Knockdowns of red Malphigian tubules reveal pigmentation roles in the milkweed bug. J Exp Zool B Mol Dev Evol 338, 382–387, doi:10.1002/jez.b.23123 (2022).

49. Green, M. M. 2010: A century of Drosophila genetics through the prism of the white gene. Genetics 184, 3–7, doi:10.1534/genetics.109.110015 (2010).

50. Khan, S. A., Reichelt, M. & Heckel, D. G. Functional analysis of the ABCs of eye color in Helicoverpa armigera with CRISPR/Cas9-induced mutations. Sci Rep 7, 40025, doi:10.1038/srep40025 (2017).

51. Socha, R. & Nemec, V. Coloration and pteridine pattern in a new, yolk body mutant of Pyrrhocoris apterus (Heteroptera: Pyrrhocoridae). Eur J Entomol 93, 525–534 (1996).

52. Melber, C. & Schmidt, G. H. Body colouration related to the deposition of pteridines in the epidermis and other organs of Dysdercus species (Insecta; Heteroptera: Pyrrhocoridae). Comp Biochem Phys A 116, 17–28, doi:Doi 10.1016/S0300-9629(96)00106-5 (1997).

53. Dorn, A. & Romer, F. Structure and function of prothoracic glands and oenocytes in embryos and last larval instars of Oncopeltus fasciatus Dallas (Insecta, Heteroptera). Cell Tissue Res 171, 331–350, doi:10.1007/BF00224658 (1976).

54. Berghammer, A. J., Klingler, M. & Wimmer, E. A. A universal marker for transgenic insects. Nature 402, 370–371, doi:10.1038/46463 (1999).

55. Grubbs, N., Haas, S., Beeman, R. W. & Lorenzen, M. D. The ABCs of eye color in Tribolium castaneum: orthologs of the Drosophila white, scarlet, and brown Genes. Genetics 199, 749–759, doi:10.1534/genetics.114.173971 (2015).

56. Lorenzen, M. D., Brown, S. J., Denell, R. E. & Beeman, R. W. Cloning and characterization of the Tribolium castaneum eye-color genes encoding tryptophan oxygenase and kynurenine 3-monooxygenase. Genetics 160, 225–234, doi:10.1093/genetics/160.1.225 (2002).

57. Kim, H. et al. A co-CRISPR strategy for efficient genome editing in Caenorhabditis elegans. Genetics 197, 1069–1080, doi:10.1534/genetics.114.166389 (2014).

58. Edgar, R. C. MUSCLE: multiple sequence alignment with high accuracy and high throughput. Nucleic Acids Res 32, 1792–1797, doi:10.1093/nar/gkh340 (2004).

59. Ronquist, F. & Huelsenbeck, J. P. MrBayes 3: Bayesian phylogenetic inference under mixed models. Bioinformatics 19, 1572–1574, doi:10.1093/bioinformatics/btg180 (2003).

60. Milne, I. et al. TOPALi v2: a rich graphical interface for evolutionary analyses of multiple alignments on HPC clusters and multi-core desktops. Bioinformatics 25, 126–127, doi:10.1093/bioinformatics/btn575 (2009).

61. Kumar, S., Stecher, G., Li, M., Knyaz, C. & Tamura, K. MEGA X: Molecular Evolutionary Genetics Analysis across Computing Platforms. Mol Biol Evol 35, 1547–1549, doi:10.1093/molbev/msy096 (2018).

62. Labun, K. et al. CHOPCHOP v3: expanding the CRISPR web toolbox beyond genome editing. Nucleic Acids Res 47, W171–W174, doi:10.1093/nar/gkz365 (2019).

63. Gloor, G. & Engels, W. Single-fly DNA preps for PCR. Drosophila Information Newsletter 1 (1991).

64. Bhattacharya, D. & Van Meir, E. G. A simple genotyping method to detect small CRISPR-Cas9 induced indels by agarose gel electrophoresis. Sci Rep 9, 4437, doi:10.1038/s41598-019-39950-4 (2019).

