## Supplementary Information for "Genome editing of the *vermilion* locus generates a visible eye color marker for *Oncopeltus fasciatus*"

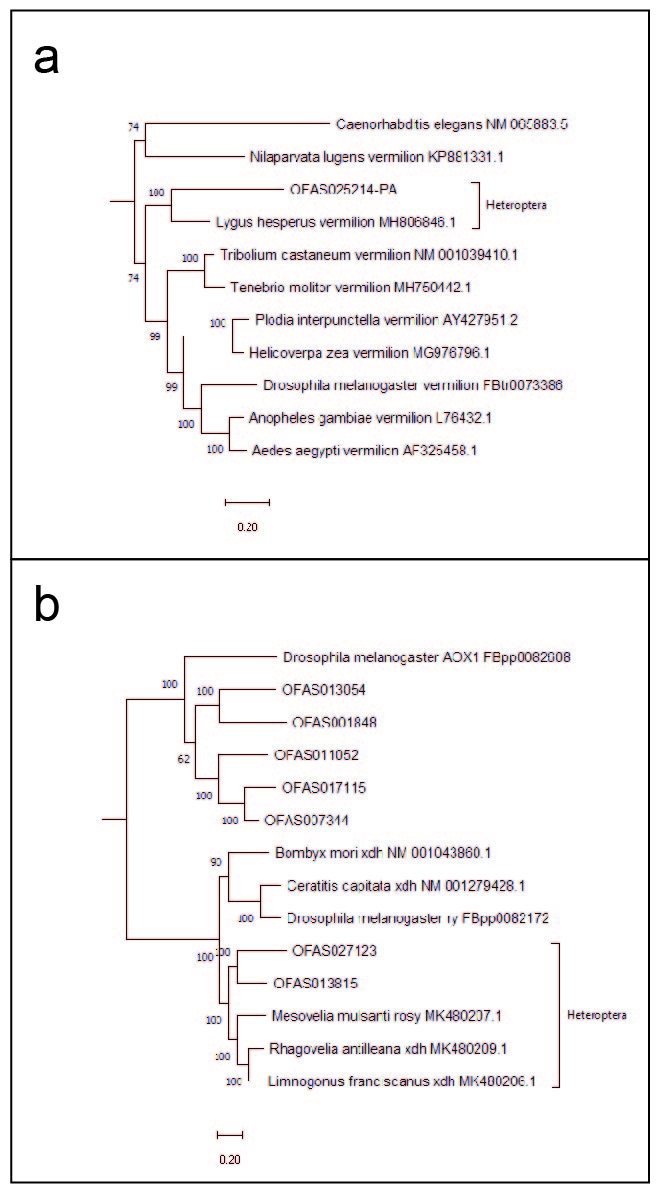

**Figure S1. Phylogenetic trees.** Phylogenetic trees of a) *vermilion* orthologs and b) *xdh/ry-*related genes. Posterior probabilities are displayed at tree nodes. a) OFAS025214 is most closely related to another heteropteran *v* ortholog, which in turn forms a clade with other insect *v* genes. b) OFAS027123 and OFAS013815 cluster with other insect *xdh* orthologs, suggesting they are both *xdh* orthologs.

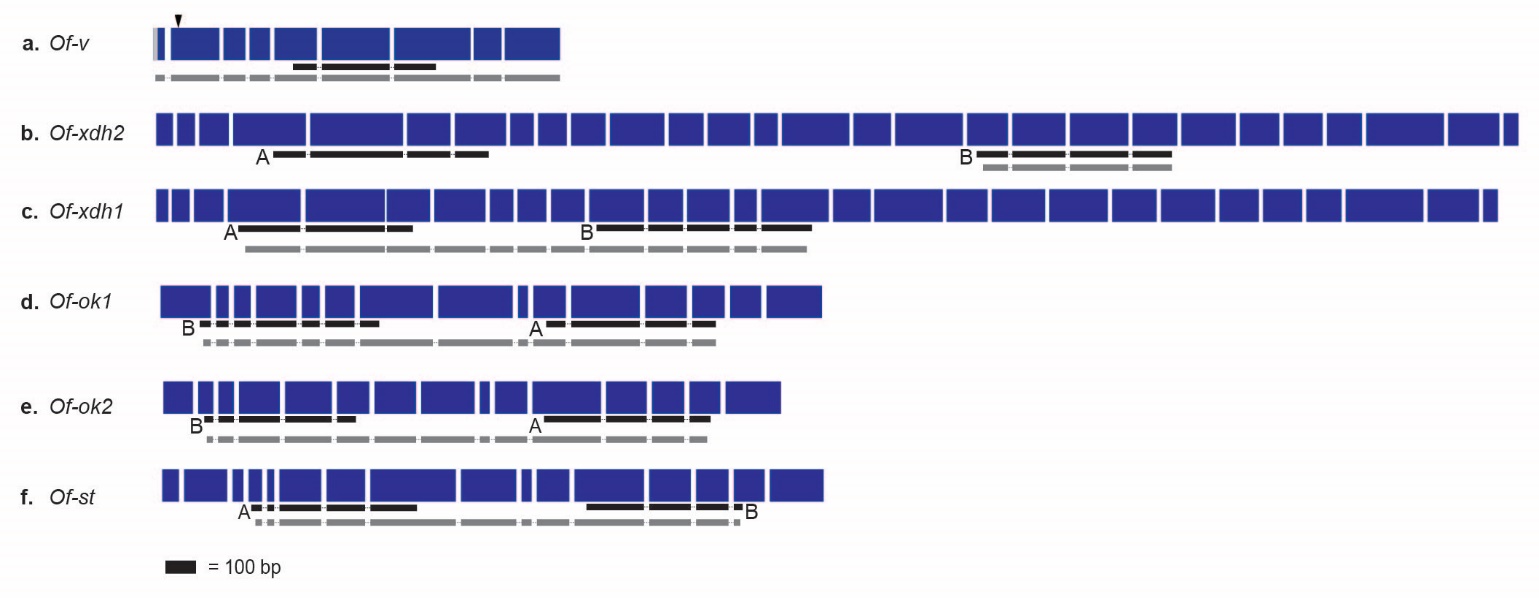

**Figure S2. Gene diagrams.** Exons of coding DNA sequence are shown in dark blue. Fragments used to synthesize dsRNAs are shown below the exons, in black. In all cases except *Of-v,* two non-overlapping dsRNAs—A and B—were used per gene. Regions shown in gray below the exons are fragments that were Sanger sequenced in the course of this study. a) *Of-v,* with partial 5’UTR shown in the first exon in light gray. The gRNA target site is in the second exon (arrowhead). b) *Of-xdh2,* c) *Of-xdh1,* d) *Of-ok1,* e) *Of-ok2,* f) *Of-st.*

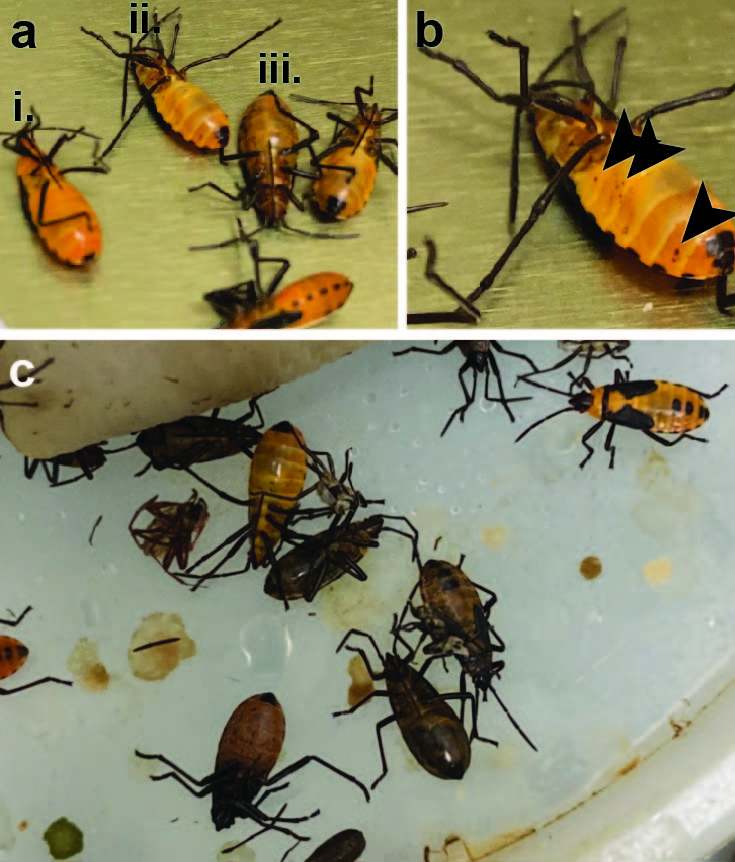

**Figure S3. Disease presentation in *O. fasciatus* population**

a) Individuals at different stages of disease. a-i) An infected individual showing signs of paralysis but not yet displaying spots of melanin; a-ii) An individual with many spots of melanin on the ventral side of the abdomen; a-iii) The entire body has darkened in an individual showing a more advanced stage of the disease. b) Close-up of individual shown in a-ii. Arrowheads point to spots of melanin. C) Dead individuals found with blackened, distended abdomens.

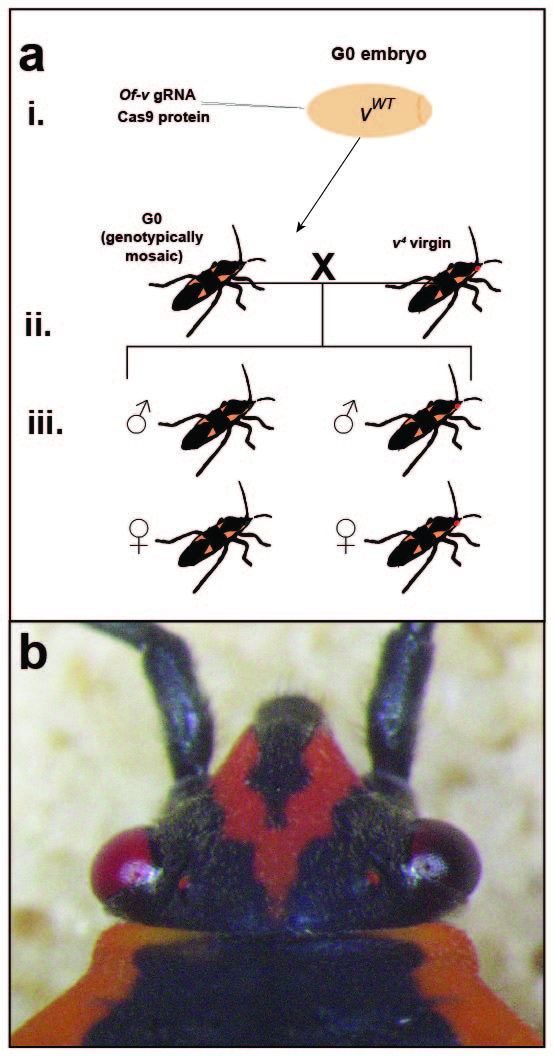

**Figure S4. Second *Of-v* CRISPR**

a) Experimental workflow for second *Of-v* CRISPR/Cas9 injections. a-i.) A gRNA targeting *Of-v* and Cas9 protein were injected into wild-type embryos. a-ii.) Each G0 individual was phenotypically scored as wild-type, having one red eye, or two red eyes before being crossed to a *v^4^* virgin of the opposite sex. a-iii.) All progeny were scored by eye-color phenotype and sex in order to calculate germline mutation rates. b) A phenotypically mosaic G0 individual with one red eye and one black eye.

**Table S1. Primer sequences**

|  | **Primer name** | **Primer sequence** |
| --- | --- | --- |
| 1 | Of-ok1-s301-FT7 (A) | taatacgactcactatagggagaGCAAGGAACCCTCAAGATCAG |
| 2 | Of-ok1-s301-RT7 (A) | aatacgactcactatagggagaGAAGGATTGAAACTGCTTCCAG |
| 3 | Of-ok2-s196-FT7 (A) | taatacgactcactatagggagaGACTGTCAGCCAGGAAG |
| 4 | Of-ok2-s196-RT7 (A) | taatacgactcactatagggagaGATATCGCCTCCAGTCC |
| 5 | Of-ok1-s301-FT7-2 (B) | taatacgactcactatagggagaGTTGAACACGACCTATTACC |
| 6 | Of-ok1-s301-RT7-2 (B) | taatacgactcactatagggagaGTTGTAACTGTCAAGTCCTGTC |
| 7 | Of-ok2-s196-FT7-2 (B) | taatacgactcactatagggagaGGAACTCTCAGTGCAGTAATG |
| 8 | Of-ok2-s196-RT7-2 (B) | taatacgactcactatagggagaGAGTAGCTGTCCAGACCAGTAG |
| 9 | Of-Xdh1-s587-FT7 (A) | taatacgactcactatagggagaGTGTATTGCATGGTCTGGCAGTG |
| 10 | Of-Xdh1-s587-RT7 (A) | taatacgactcactatagggagaGAACCTCTCTGAGTGTTACAGGTC |
| 11 | Of-Xdh2-s1491-FT7 (A) | taatacgactcactatagggagaGTCTCAGTGTGGCTTCTGTACTCC |
| 12 | Of-Xdh2-s1491-RT7 (A) | taatacgactcactatagggagaCAGCTCCAATCCTCAGGCCATTAC |
| 13 | Of-Xdh1-s587-FT7-2 (B) | taatacgactcactatagggagaGCAATCCAGGAGGAGAGATG |
| 14 | Of-Xdh1-s587-RT7-2 (B) | taatacgactcactatagggagaCCTGGAGTAATATCCTTGGC |
| 15 | Of-Xdh2-s1491-FT7-2 (B) | taatacgactcactatagggagaGTAATCGTGTTGTGAGCAGAG |
| 16 | Of-Xdh2-s1491-RT7-2 (B) | taatacgactcactatagggagaCAGCTTGACTCTGGAGTG |
| 17 | Of-st-FT7 (A) | taatacgactcactatagggagaGAATCAGGACAACTGGTAGC |
| 18 | Of-st-RT7 (A) | taatacgactcactatagggagaCGCATTCCAAGATGCCATC |
| 19 | Of-st-FT7-B (B) | taatacgactcactatagggagaGTGAACAGAGATCAAGAAGG |
| 20 | Of-st-RT7-B (B) | taatacgactcactatagggagaGCAGCCAAGATATTACATCAG |
| 21 | Of-ver-RNAi-FT7 | taatacgactcactatagggagaGAGACTACCTCTGCCCTGCTTC |
| 22 | Of-ver-RNAi-RT7 | taatacgactcactatagggagaGCTTTGTGTGTAAATCTCCTCTCACCTC |
| 23 | gRNArev | AAAAGCACCGACTCGGTGCC |
| 24 | Of-v-gRNA-1 | ttaatacgactcactataggTCTTCCGAGATCCAGAAATGgttttagagctagaaatag |
| 25 | Of-v-F | CACCAGGTACACTATAGTCGTAGCG |
| 26 | Of-v-R | GCTCACTTTATCTTGTAAACTGCCTTCATTG |
| 27 | Turbo GFP 1R_T7_2 | ttaatacgactcactataggCTCGGTGTTGCTGTGATCCTC |
| 28 | T7-tGFP-F | taatacgactcactatagggagaATGGAGAGCGACGAGAGC |
| 29 | AmpR-gRNA-A | ttaatacgactcactataggTAGATAACTACGATACGGGgttttagagctagaaatag |
| 30 | Of-v-fullCDS-F | CAGTGCAGAGAGCGTCTTC |
| 31 | Of-v-fullCDS-R | CAAAACATTTTTAATCTTCTTCGTGATCTTC |

**Supplemental file: Sequences**

>Of-v-consensus

CAGTGCAGAGAGCGTCTTCGATGGAGTCCGACAGCAATTGCAGATCTTCCGAGATCCAGAAATGTGGAGAGCAAAGTGGAATGCTGTATGGGGATTACCTCCAACTTGACAAAATACTGAAGGCCCAAAGATTGCTGAGCGAATCCAACAACGCCACTGTAATGGACGAACATTTATTTATCGTCACACATCAAGCTTACGAGCTATGGTTTAAACAAATTATATACGATTTAGATTGGGTCCGAGACGTCTTTAGCAATCATAAGGGACTTAATGAGCGACAAACTCTGGATGTTGTCAAACGATTAAATAGAATTGTATTAATCCTAAAGTTGCTAGTGGAACAAGTAATGATACTTGAAACAATGACTCCATTGGACTTCATGGAGTTTAGAGACTACCTCTGCCCTGCTTCTGGTTTCCAAAGCGCACAATTTCGCCTCATTGAGAACAAACTTGGAGTAAAAACGGAACATCGGGTCCGTTCCTTTCAACATTACTCTACTGTTTTTGAGAGAGACAAGGAAGCACTTAATGAAATCAAGAAATCAGAAACCGAACAGTCTTTAGCAGATCTGATTGAGCAGTGGCTTGAGCGTACGCCTGGCCTGGATGAAAATGGATTCGATTTTTGGGGAAAATATATAAAAAATGTTGAGTTTCTCCTTCAGCATTATTATGAAAAGGCAATGAAAAGCAATTCAGAAACAGAGAAAACTACAAAGTTAGCAGAATATGAAAAGAAGAAAACTATTTTTAATTCAGTTTTAAATAAAGATATGCATGACTTATTTGTACAAAGAGGTGAGAGGAGATTTACACACAAAGCATTTCAAGGTGCCATTATGATAACATTTTATAGAGATGAGCCCCGTTTCAGCCAACCTCACCAAATTCTTACTTTATTAATGGATATCGATTCCCTAGTCACCAAATGGAGATATAATCATGTAATTATGGTCCAAAGGATGATTGGGTCTCAAAGTGGAACAGGAGGTTCATCAGGTTATCACTACTTGCGCTCAACTCTTAGTGATAGGTATAAATGTTTTTTGGACTTGTTCAACCTTTCATCATTTTTATTGCCTCGAAATTTCATCCCCCCATTAAGCTCATCAGTAAGGAATCAAATTAATATACACACCTGGAATACAAAGAAACAAACAGAAGATCACGAAGAAGATTAAAAATGTTTTG

>Of-ok1-consensus

AGTGGTGTTGCAGAACCAGGATCATTGATGGCCATTATGGGTCCTAGTGGAGCAGGAAAAACAACGTTACTGGCAACAATTAGTCAAAGGCTCAAAGGTGATGTAAAAGGTAAAATTTTAGTCAATGGACAACAAATCTCAAGAGAATTAATGATGAGGGTATCAGCTTTCCTTCCTCAGCAAGATCTGACTATTGATAGTCTAACAGTTTATGAACATCTACAATTAATGGCTAGATTAAAATTAGACAGAGATATAAGTCCTGCTGAAAGGACTGAGAGAATTCTAGCTATTTTAACTGAGTTGGGTATCAATAAATGCAAACACACATATATTAAATCTCTCTCAGGTGGTGAAAGACGCCGAGTATCTTTAGCTGTTGAGTTGCTTACAGATCCAGCCATTCTCTTCTGTGATGAGCCTACGACAGGACTTGACAGTTACAACGCTACATCTGTGATAGAGCAGCTTCGAAGGCTTGCTTCCACAGGGAAAGTTATTATATGTACAATACACCAGCCAACTTCAGGAATATTTGAATTGTTTCAGACTGTTTATTTGCTTGCTTCTGGGGGCAAACTTGCCCTTACTTGTCCTATTCACGAAGCTGCATTGTTTTTCCAAAAAATGGGTATGGTTTGTCCTCCTACATATAACATAGCTGAATTTTTAGTCAGCCAACTTGCACATCAAGAAGATTCTGAAAGTGAATTGAGACTTTTTAAAATTTGTAATGAGTTTAAGTTGAGTGAACATTATGAAAATCTTATGAAAAAATTAGATGGAGAATTGAATCATGCAAAGTTTATGGTTCCTAATTATTCAACTAATGCACAAGATTTTAAACATTCTTTTCAGTCTTTATATGAGTATGGAGATGAGTTTCACAAGTACATGTCTATCAAACGTCCTTCTAGACTTACTCAGTTAGGATGGCTAATGTGGAGGTCTTTCCTTGACATTGCAAGGAACCCTCAAGATCAGTTAATTCGTCTTGCTTTTTTTATGTTTATGGCTATTCTCATATCAACTCCTTATGTTGGGCTCAAAGTAGATCAAGAAGGAATTCAAAATTTGCAAGGTCTGCTTTACTTAATAATTGTTGAGACAATATTCACTTATACATACAGTGTGAGTCATACCTTTCCTTCCGAAATACCAATATTGCTGCGTGAAGTCAACAATGGCTTGTACACTCCTGGACCTTATTATGTTTCAAAGATGATAATTTTGCTTCCTAGAGCTCTTTTTGAGCCAATAATCTATGCAGCTGTGGTGTTTTGGATACCTGGTTTGTTAGGAGGTTTTGCTGGATATCTTGAATTTTGCATACCTATTATTGTTTGTTCAATGGCTGCAACTGCTTATGGGTGCGTTATTTCTGCTATATTTGAAGATATTTCTACTGGTTCTCTATTGTCTGTGCCATTTGAACAGTTTTGTCTACTTTTCTGTGGATTGTATTTGTCATTGAGTGATGTACCATTTCATCTCGCATGGGTGAAGTACATATCAATATTTTNCTATGGG

>Of-ok2-consensus

GGTCCCAGTGGTGCAGGTAAAACAACTTTGCTAGCGGCAATAAGCCAGAGATTTTCTGGAGAGCTGAGGGGCGAGATCCACCTTAGCGGACGGCCGGTGGACAAGGAGCTCATGATGAAAATGTCTGGCTTTGTCCCACAGCACGACTTGGCCGTCAAGACCCTCACTGTTAGCGAACATTTAAATTTCATGGCCGTGCTGAAGATGGATCGGCGGGTCTCGAAACTCCAGCGGCGCCGGATAGTGGAGTCTCTTCTGCTAGAGCTGGGCCTGAATGACAGCGCAGACTCCAGACTCGGCGCGCTCTCTGGAGGGGAGAGGAAGCGGCTCAGCCTCGCTGTTCAGCTGGTGACAGATCCTCCGGTCCTCCTGTGTGACGAGCCCACTACTGGTCTGGACAGCTACTCGGCTGGCAGTGTGGTCAGCCTGCTCCGACAACTTGCTTCCAGAGGTAAAGCTGTACTGGCGTCTGTCCACCAACCAGCCTCTGGGATCTTCGAACAGTTCGACACGGTGACTCTCCTTGTTCCCGGTGGTCGGATGGCCTACTTCGGCGAGGTGAACTCCGCCAAGCTGTACTTTTCGCAGATGGGTCTGAACTGTCCGACCTCCTACAACACTGCCGAGTTCCTGGTGACCCAGCTGAACACCCTCCCAGACAAGCTGACGGAGAAGTTCGTCACCACGAACACCTTCGTCACCATGCACAAGGATATCGAGACAGTCAAGAAGAAGACCTTGAACGACACCCTCATCTATGGAATGGAAGAAAAGTTTTTGAAGTTTTATTCAGTGCAACCATCGACGGATAAGACTCAGATAAAGTGGCTACTGTGGCGTTCGGCCCTCGACATGGTTCGAGACTCGCACAGGCTATTCGTCAGGCTCATCATGTATCTCGTGACGGCGATATTGATATCCACCCCGTTCGTGGGGACGACTGTCAGCCAGGAAGGCATCCAGAACGTCCAGGGTCTGAGCTACTCCGTCATCACGGAGACAGTGTTCTCCCACGCGTACGCAGTGATGCATACCTTCCCCTCCGAAATCCCCATACTCCTGAGGGAGGTCAGCAATGGGGTGTACAAACCCGCCCCGTACTACCTCTCCAAAGTCCTCTTCCAGATTCCGAGGACAATCATCGAGACAATTCTATTCTGCTTAGTGATATACGTGATAGCTGGAAAAGGCTTAGAAGTGAATTTCTTCTACTTCTGTATTCCAGTGGTCGTCGTGGCAGTAGCTTCCACAGCCTACGGCTACTGCATTTCGGCCATGTTTGAGAGTATCAGTACAGCGTCCTTGTTGTCTGTGCCTGTAGATTTCATCTCCTACACGTTCAGTGGGATTTTTCTTCAGCTCAGCACGGTTCCTCTCTACCTCTCCTGGGTGAAGT

>Of-st-consensus

GGAGGAATGTTAGTTGATGGTGATGTAAGACTTAATGGAAAGAAAATGGATTCATCTATGTGCAAAATGTCTGGACTTATGAGGCAAGAAGATCTTTTTATCGGCGACCTCACAGTGAAGGAACATCTCATGTTTATGGCAAGGCTCAAATTAGACAGAAGATTCAATTCTGTTGAAAGAGAAAATCGAGTATGTTCCTTGATAAAGAATCTTGGACTGAGCAAGTGTGCTCATCGAAAAATTGAAGGCAGCACTTTTGGCAATGTCACAGCATTGTCTGGTGGAGAAAAAAAACGTCTATCATTTGCTACTGCATGTCTCCCAGATCCTCCTCTACTCTTCTGTGATGAACCAACAACAGGTCTTGATTCATTAAATGCAATAAGAATTGTCCGAATGATGGCATCTTGGAATGCGACAATTTTGTGCACCATTCATCAGCCAAGCCCTGAAATTTTGAAGTATTTTTCTTCTGTTATTTTAGTTGCTGATGGTAGCCTTGTATTTTCGGGTACAGTTGAGAATGCTGTCGATTTTTTAAAAAGTGCTGGTTATGAACAAAATGAACACAAAAGTGCTGGAGAATTTCTTGTGACATGTTTAGCTCCAACCCCTGGGTCTGAACAGTCCACTAAAATCGCTGTGAGAAGACTAGCACATCAGTTTGCTCTTTCAGAGTACGCAAAAACCAGTGAAATGCACATACAATTGCACAAACAAACTGAAAGGAAGACAAAAAATCGCGATCTTCATGATCATAAATCTTCATTTTGGATGTGCAAATTTTTCTGGCTTCTAATTAGAGGAATAATAGAAATTCTTCGAAACCCACAAATACAATGGGTAAAAATAGGACAAAAAGTTGTAGTAGCCCTGATGGCAGGCCTGTGTTTTGTTGGAGCTGTGAACAGAGATCAAGAAGGTCTGCAAGCTGTTCAGGGTGCCCTGTTTATTCTTGTAGCTGAGAACACTTTTTCTCCCATGTATTCAGCACTTGCACTCCTTCCCAAAGAATTTCCACTTCTCCTCCGGGAACTCCGAGATGGAGATTACCATCCATCAGCATTGTACCTTGCTAAAATAATATCATATATTCCAGGACTTGCATTTGAGGCTGTGCTGTTTACTGCTATTGTATATTGGCTAGCTAATTTAAGGGATACCGTGTTTGCATTTTCCATGGTACTGTTATGTAATATTTTATGTGTATTGGTTTCATCAGCTTGTGGTATGATGTTTGCTAGTATTTTTGACAATTATGAGCTGGCAAATATTTATTTGATTCCATTTGATAATGCTATGATGATTTTGTCTGGGGTATATAT

>Of-xdh1-consensus

AGGGCATAGGGAGTTTTGCTGATAAACTTCATCCTGTCCAAGAGAGGATCGCCAAGGCACATGGTTCGCAATGTGGCTTCTGCACTCCTGGCATCGTTATGTCCATGTACGCACTCCTCAGGTCTCTCGACAGGAAACCTACAATGGATGACATTCAGACTGCTTTCCAAGGGAACTTGTGTCGCTGCACAGGATATAGACCAATTCTTGATGGCTACAGGACGTTCACCGGTCCGCCTCTGGAGTACAAAGGCAATTCCTCTTGTTCAATGGGAAACAAATGCTGTAAAGTCAATGGCCATATTGATAAAAGCATTGATGATAGGGGAAATAACAGTGTAGAATCTGAGGTCCCGGTCATTCTCTTCAACGGGGAAAAGCTGAGCAAATATGATCCAACTCAAGAAATAATATTTCCACCTGAACTTGCTATTTCAGACAAATACGACAAGGAGTCCTTACACATCAAAGGACCAAGGATAGCTTGTTATAGACCTGTAACACTCAGAGAGGTTCTTGAATTGAAGGCTTTACATCAAGATTCAGTTATTATCAATGGAAACACTGAAGTTGGAATTGAAATAAAATTTAAGAACATTCTCTATTCTCATGTGATAGTGGCAAATATGGTCTCTGAGCTATCACAAATATCCTTGGAAGAGGGTGGTGTAAAGATTGGTGCAGCTGCAACTATTAATGAAATTGAAGAATTTTTGATAGAAACACTAGCTTTGAAAGAAGAAGCTGCTTGCAGAATATTTATATCTATGCTTTCTATGATCAAATATTTTGCTGGAAAACAAATAAGAAATGTTGCTGCAATTGGAGGGAATATAATGACTGGTAGCCCTATTTCTGATCTGAATCCTATTTTTATGGCTGCTGGAGCTGAACTAGAATTTCAAACTACTGGAGGTTCGCGTAAGTTGATAATGGATAACAACTTCTGGATTGGTTATCGAAAAAATATTGTACAAAAGAATGAAGTACTGACCTCCATAGTTCTACCTTTCACTTCTAAGAACCAATATTTTAAGGCATACAAGCAATCCAGGAGGAGAGATGACGATATTGCAATTGTAAATGCAGCATTCTTTTTGGACCTTGAAAGAAATACAATTAAAAATATTCATATGGCTTTTGGTGGAATGGGACCAACTACTCTTATGGCCATCAGAACTGAAAAAATGTTAATTGGGAAGAAATGGAATGAATCAACATTGAATGAAGCAATGTCCTCATTGACAGTCGAGCTTTCATTACCTTCTGAAGCCCCAGGTGGAATGGTGCAGTTCAGGAAATCTCTTACCCTCAGCTTTTTCTTCAAATTCTACCTTTATGTTGCCAATGAAATGGAAAAAATGGGCTTAAATTATTCTGTAGACAATGAACTTAAAAGTGCCATTGATGAAATGCATATTTTAAGCCCCACTAGTTCACAATTTTTTTCTATTAATCCTGCAAAGGAGGAGTACAGCACTGTTGGCCAACCCATAGTGCATCAATCAGCATTCAAACATGCTACTGGGGAAGCTATTTATTGTGATGATATGCCCAAACTTGAAAATGAAGCCTACTTAGCTCTTGTACTAAGTAGCAAACCCCATGCTAAGATTCTCAGCATTGATAGTTCAGAAGCTCTGAAATTGGAAGGAGTG

>Of-Xdh2-partial

AAAGGAGTCCAAGTATAGTGCTCTTGCACTTCCTGTTGCTGTTGCTGCTTCAAAACTAAATCGTCCTGTAAGAGCTATGCTGGACCGAGATGAAGATATGATGATAACTGGACATAGACACCCTTTCTTAGGGAAATACAAAGTTGCTGTTACAAAAGAAGGGAAAATGATAGCTTGTGAAGTTAAAATTTATTGTAATGCAGGGTTTACATTGGATCTTTCTTGTTCTGTTATGGATAGGGCAATTTTCCACTTTTCAAATGCCTTCTTTATTCCAAATGTAGTAGTCTATGGTTATTGCTGTAAGACCAATATCCCTTCAAATACTGCTTTCCGAGGATTTGGAGGGCCTCAGGGTATGTTCATAGGCGAGACTATGGTCACTCATATTGCCGAAACTCTCAGCCTAGACTCTCAGCAGGTTAGAGAAATAAACTTCTACAAAGAAGGATATATCACCCACTATAACCAAGAATTATCATATTGCACCTTAGATCGCTGTTGGAATGAGTGTATTTCTAAATCAGATTACCACTCCAGAGTCAAGCT

>Of-exon2-WT

TAGTCGTAGCGCTACCATAAGTAAGACTTAAATACTACTTGCTCGAGGAAGAGGCTTGAAGGAAATCCATCGGGAAGGGAGATCGCATGAAAGCTTGATTTTCTTTCAAATGTAAATGTACGTGTTCTGTTTCAGTTGCAGATCTTCCGAGATCCAGAAATGTGGAGAGCAAAGTGGAATGCTGTATGGGGATTACCTCCAACTTGACAAAATACTGAAGGCCCAAAGATTGCTGAGCGAATCCAACAACGCCACTGTAATGGACGAACATTTATTTATCGTCACACATCAAGGTAATAAACTCAACTCCACTCAAGAAATTTACTTCTAAAATCAATATGCTGAAAATAATTGATCAAATGCAGGTTACTGTTTGTATAAGCAGCTGAAAAGGCACCTCTTTTGCATTCGGTCGGTTTGCTATTTACCCATTGATAAACATTGCATCCTTTGGCAGAAACATCTTGGTTTATACAACTAACTATAAATTCTATGTTGATTAACC

>Of-exon2-v1

CTATAGTCGTAGCGCTACCATAAGTAAGACTTAAATACTACTTGCTCGAGGAAGAGGCTTGAAGGAAATCCATCGGGAAGGGAGATCGCATGAAAGCTTGATTTTCTTTCAAATGTAAATGTACGTGTTCTGTTTCAGTTGCAGATCTTCCGAGATCCAGAAGATGTGGAGAGCAAAGTGGAATGCTGTATGGGGATTACCTCCAACTTGACAAAATACTGAAGGCCCAAAGATTGCTGAGCGAATCCAACAACGCCACTGTAATGGACGAACATTTATTTATCGTCACACATCAAGGTAATAAACTCAACTCCACTCAAGAAATTTACTTCTAAAATCAATATGCTGAAAATAATTGATCAAATGCAGGTTACTGTTTGTATAAGCAGCTGAAAAGGCACCTCTTTTGCATTCGGTCGGTTTGCTATTTACCCATTGATAAACATTGCATCCTTTGGCAGAAACATCTTGGTTTATACAACTAACTATAAATTCTATGTTGATTAAC

>Of-exon2-v3

TATAGTCGTAGCGCTACCATAAGTAAGACTTAAATACTACTTGCTCGAGGAAGAGGCTTGAAGGAAATCCATCGGGAAGGGAGATCGCATGAAAGCTTGAGTTTCTTTCAAATGTAAATGTACGTGTTCTGTTTCAGTTGCAGATCTTCCGAGATCCAGAGCAAAGTGGAATGCTGTATGGGGATTACCTCCAACTTGACAAAATACTGAAGGCCCAAAGATTGCTGAGCGAATCCAACAACGCCACTGTAATGGACGAACATTTATTTATCGTCACACATCAAGGTAATAAACTCAACTCCACTCAAGAAATTTACTTCTAAAATCAATATGCTGAAAATAATTGATCAAATGCAGGTTACTGTTTGTATAAGCAGCTGAAAAGGCACCTCTTTTGCATTCGGTCGGTTTGCTATTTACCCATTGATAAACATTGCATCCTTTGGCAGAAACATCTTGGTTTATACAACTAACTATAAATTCTATGTTGATTAAC

>Of-exon2-v4

TAGTCGTAGCGCTACCATAAGTAAGACTTAAATACTACTTGCTCGAGGAAGAGGCTTGAAGGAAATCCATCGGGAAGGGAGATCGCATGAAAGCTTGATTTTCTTTCAAATGTAAATGTACGTGTTCTGTTTCAGTTGCAGATCTTCCGAGATCCAGATCTCCTAATGTGGAGAGCAAAGTGGAATGCTGTATGGGGATTACCTCCAACTTGACAAAATACTGAAGGCCCAAAGATTGCTGAGCGAATCCAACAACGCCACTGTAATGGACGAACATTTATTTATCGTCACACATCAAGGTAATAAACTCAACTCCACTCAAGAAATTTACTTCTAAAATCAATATGCTGAAAATAATTGATCAAATGCAGGTTACTGTTTGTATAAGCAGCTGAAAAGGCACCTCTTTTGCATTCGGTCGGTTTGCTATTTACCCATTGATAAACATTGCATCCTTTGGCAGAAACATCTTGGTTTATACAACTAACTATAAATTCTATGTTGATTAAC
